# High-resolution whole-brain magnetic resonance spectroscopic imaging in youth at risk for psychosis

**DOI:** 10.1101/2025.06.22.660965

**Authors:** Edgar Céléreau, Federico Lucchetti, Yasser Alemán-Gómez, Daniella Dwir, Martine Cleusix, Jean-Baptiste Ledoux, Raoul Jenni, Caroline Conchon, Meritxell Bach-Cuadra, Zoé Schilliger, Alessandra Solida, Marco Armando, Kerstin Jessica Plessen, Patric Hagmann, Philippe Conus, Antoine Klauser, Paul Klauser

**Affiliations:** Centre for Psychiatric Neuroscience, Department of Psychiatry, Lausanne University Hospital and University of Lausanne, Lausanne, Switzerland; Service of Diagnostic and Interventional Radiology, Department of Medical Radiology, Lausanne University Hospital (CHUV), Switzerland; CIBM Center for BioMedical Imaging, Lausanne, Switzerland; Division of child and adolescent psychiatry, Department of Psychiatry, Lausanne University Hospital and University of Lausanne, Lausanne, Switzerland; Centre Neuchatelois de Psychiatrie, Neuchatel, Switzerland; Service of General Psychiatry, Department of Psychiatry, Lausanne University Hospital and University of Lausanne, Lausanne, Switzerland; Swiss Innovation Hub, Siemens Healthineers International AG, Lausanne, Switzerland

**Author notes:** Corresponding author: Paul Klauser; Centre de Neurosciences Psychiatriques, Route de Cery 11bis, 1008 Prilly, Switzerland.

**Keywords:** Magnetic Resonance Spectroscopic Imaging, Clinical High Risk, Schizotypal Personality Disorder, N-Acetylasparate, Inositol, Voxel-based Analyses

## Abstract

Advances in three-dimensional magnetic resonance spectroscopic imaging (3D-MRSI) allow for the high-resolution mapping of multiple neurometabolites throughout the entire brain in vivo and within clinically compatible time frames. Leveraging this capability, we created a voxel-based pipeline that corrects and spatially normalizes whole-brain maps of total N-acetylaspartate (tNAA), myo-inositol (Ins), choline compounds, glutamate + glutamine, and creatine + phosphocreatine. We examined 2 different 3D-MRSI dataset: first, a clinical sample of adolescents and young adults at risk for psychosis (n= 21) meeting DSM-5 criteria for Attenuated Psychosis Syndrome (APS) or Schizotypal Personality Disorder (SCZT), and age-/sex-matched healthy controls (n =13); and second, a non-clinical sample of adolescents (n = 61) scanned on a different site. The objective of the study was threefold: first, to assess the reproducibility of 3D-MRSI measures across datasets and scanning sites; second, to validate the feasibility of whole-brain, voxel-based analyses on 3D-MRSI data; and third, to test the sensitivity of this approach. Metabolite distributions showed reproducible regional variation in standard space between the two independent samples and scanning sites (r ranging from 0.82 to 0.99). Relative to controls, at-risk participants exhibited higher tNAA levels in frontal grey matter; the SCZT subgroup additionally displayed widespread cortical and subcortical elevations of Ins levels compared with both APS and controls. Voxel-based analyses of structural (i.e., gray and white matter volumes or densities) and diffusion (i.e., generalized fractional anisotropy) parameters yielded no significant differences between patients and controls. These preliminary findings suggest that high-resolution 3D-MRSI may be sensitive enough to detect subtle neurometabolic alterations at the group level in the early stages of psychotic disorders when structural or diffusion measures show no difference. High-resolution whole-brain metabolic mapping may have the potential to help with early identification of young people at risk for psychosis or other mental disorders.

## 1. Introduction

Magnetic Resonance Imaging (MRI) has played an important role in the search for biomarkers in psychotic disorders, including schizophrenia (SZ). Structural and diffusion brain MRI allow the detection of group-level differences in cortical and subcortical regions of patients with established SZ, including widespread cortical thinning, loss of volume in the hippocampus, amygdala, and thalamus as well as altered diffusion properties in the vast majority of white matter tracts (for the ENIGMA Schizophrenia Working Group et al., 2016; Kelly et al., 2018; P. Klauser et al., 2016; Van Erp et al., 2018).

However, during the earliest stages of the disorder, when structural alterations are less important, detecting differences between youths at risk for psychosis and healthy controls becomes more difficult, and the effect sizes remain small, even in meta-analyses with large samples (ENIGMA Clinical High Risk for Psychosis Working Group et al., 2021). This could be due to sensitivity limitations of the current techniques that only detect the water signal. Advances were achieved in the 1990s with Magnetic Resonance Spectroscopy (MRS), a technique that allows the detection of multiple metabolites in a single volume of interest (SVS-MRS) (Van Der Graaf, 2010; Zhu & Barker, 2011). At the same time Magnetic Resonance Spectroscopic Imaging (MRSI) was developed with the goal to combine spatial with spectroscopic information (Bogner et al., 2021; Maudsley et al., 2021).

Although MRSI was first proposed in the 1980s, applications were limited for an extended period due to lengthy acquisition times and coarse spatial resolution. Subsequent progress addressed two distinct challenges: (i) water and lipid signal handling and (ii) sampling/reconstruction efficiency. Concerning the first point, approaches such as outer-volume suppression and inversion-recovery lipid nulling were initially employed (Tkáč et al., 2021). More recently, advances in post-processing, including lipid suppression and low-rank reconstructions have enabled reliable, near-whole-brain coverage (A. Klauser et al., 2019; Maudsley et al., 2009). Regarding the second point, a broad family of acceleration and encoding strategies has been developed: parallel imaging (SENSE/GRAPPA), compressed sensing (CS), and low-rank modeling combined with fast k-space trajectories, including echo-planar spectroscopic imaging (EPSI/PEPSI) (Zhu & Barker, 2011), spirals, concentric rings (Esmaeili et al., 2021; Hingerl et al., 2018). Moreover, subspace-based methods, such as SPICE (spectroscopic imaging by exploiting spatiospectral correlations), have improved the balance between spatial coverage, spectral resolution, and scan time (Lam et al., 2016). Recently, rosette-based MRSI demonstrated its acceleration potential and reproducibility (Z. Huang et al., 2025). Finally, ECCENTRIC was developed as a fast approach for high-resolution MRSI to improve spatial response function and spectral quality (A. Klauser et al., 2024a). Meanwhile, free-induction-decay (FID) MRSI with a very short TE emerged as an acquisition strategy that samples immediately after excitation. This reduces T2 and J-coupling effects while enhancing the signal-to-noise ratio (SNR) and metabolite detectability (Hangel et al., 2021a; Moser et al., 2020). Combining FID-MRSI with the aforementioned acceleration and reconstruction approaches enables high-resolution whole-brain mapping at 3T in clinically practical times (A. Klauser et al., 2022). A recent review provides a comprehensive overview of these developments (Bogner et al., 2021). In this study, we used a whole-brain ^1^H-CS-FID-MRSI acquisition, referred to as 3D-MRSI.

Fast 3D-MRSI technique at 3T scanners can resolve reliably 5 metabolites over the whole brain, some of which are composite signals comprising multiple metabolites due to overlapping spectral peaks. The most prominent signal is N-acetylaspartate (NAA), detected with N-acetylaspartyl glutamate (NAAG), which, together, forms the total NAA (tNAA) composite signal. tNAA is generally considered a marker of neuronal viability, synthesized in neuronal mitochondria, and reduced in case of neuronal loss such as in Alzheimer’s disease (Jessen et al., 2001; Van Der Graaf, 2010). Creatine and phosphocreatine, both acquired as total creatine (tCr), play a crucial role in brain energy metabolism. It was previously largely used as a normalization tool for other metabolites, although age and some diseases seem to alter its concentration (Minati et al., 2010; Schuff et al., 2001). Choline-containing compounds (Cho) play an important role in membrane synthesis and thus have been widely used for brain tumor characterization (Herminghaus et al., 2002). Myo-inositol (Ins) is a sugar involved in osmotic balance and mainly found in glial cells, especially astrocytes. Elevated levels of Ins have been previously associated with glial proliferation and reported in various neurodegenerative disorders, including using MRSI (Hu et al., 2024; Kim et al., 2005; McKiernan et al., 2023). Finally, glutamate (Glu), the most abundant excitatory neurotransmitter in the brain, and glutamine (Gln), which is both a precursor for Glu synthesis and a product of Glu metabolism in the Glu-Gln cycle, are both being identified as a single contribution (Glx) due to strong signal overlap. The relative difficulty to resolve Glx has led to mixed results in most brain pathologies (Poels et al., 2014; Ramadan et al., 2013).

As detailed before, 3D-MRSI addresses both the detection of 5 signals other than water (the limitation of structural and diffusion MRI) and can map them on most of the brain volume (the limitation of classic SVS-MRS). Thus, 3D-MRSI is a promising approach for detecting subtle changes during the earliest stages of psychiatric disorders such as psychosis. People with psychosis, including schizophrenia, suffer from various disabling psychiatric symptoms (e.g., hallucinations, delusions, social withdrawal, cognitive impairments) that often lead to a poor quality of life, multiple hospitalizations and long pharmacological treatments (Tandon et al., 2008). A prodromal phase generally precedes the first psychotic episode. During this stage, patients at-risk for psychosis experience a deterioration in cognition (Catalan et al., 2021) and global functioning (Oliver et al., 2020), as well as the onset of subclinical psychotic symptoms (e.g., hallucinations with insight) (Fusar-Poli et al., 2013). The rate of conversion from the prodromal phase to first episode psychosis is around 20% within 2 years, but most of these patients will develop at least one psychiatric disorder later in life (Clark et al., 2016; Salazar De Pablo et al., 2021). Therefore, patients at risk for psychosis are an important group to target for early detection and intervention.

Despite some residual controversy, the international psychiatric classification DSM-5 (*Diagnostic and Statistical Manual of Mental Disorders*, 2013) identifies these patients in two categories. First, the schizotypal personality disorder (SCZT) which includes odd beliefs and thinking, suspiciousness and ideas of reference, constricted affects, and social withdrawal (Kirchner et al., 2018). The DSM-5 also includes in its appendix the Attenuated psychosis syndrome (APS) which comprises attenuated delusions, hallucinations and disorganized speech for at least a week in the past month and that have appeared or worsened in the last year (Salazar De Pablo et al., 2020). APS is a partial transcription of the “clinical high-risk (CHR)”, “at-risk mental state (ARMS)” or “ultra-high risk (UHR)”, states that have been widely documented in the psychiatric literature.

Previous research utilizing SVS-MRS and MRSI has demonstrated alterations in metabolic profiles among individuals with psychosis. In those with SZ, meta-analyses report reduced tNAA across various cerebral regions (Iwata et al., 2018; Yang et al., 2023), elevated Cho within the basal ganglia (Yang et al., 2023), increased Glx in the medial prefrontal cortex (MPFC) of unmedicated patients and in basal ganglia (Plitman et al., 2019; Poels et al., 2014), as well as heightened Ins in the MPFC (Das et al., 2018). In contrast, recent studies using ultra-high field imaging (i.e., 7T), showed reduced Glu and GABA levels in the insula of patients with first episode psychosis, followed by a normalization of Glu levels in remitters (Sonnenschein et al., 2022), as well as a reduction of Glu/GABA ratio in the dorsolateral prefrontal cortex, which is already present in CHR (Mayeli et al., 2022).

In CHR patients, meta-analyses show elevated Glx, tCr and Ins in the prefrontal cortex, and decreased Glx and tCr in the thalamus (Romeo et al., 2020; Wang et al., 2020). A recent 7T MRSI study, also reported increased GABA/Glu ratio in the thalamus of CHR individuals, but this finding was not replicated in a recent paper (Keihani et al., 2025). Most of these prior investigations, including those employing MRSI, examined only specific brain regions rather than using a whole-brain approaches. Nevertheless, a recent MRSI study took a whole-brain approach to compare individuals with SZ, bipolar disorder and healthy controls. This study identified increased levels of Ins, tNAA, Cho and Glx in SZ patients, predominantly in the cingulate gyrus and frontal brain regions (Bustillo et al., 2021). Whole-brain MRSI approaches have also been used outside of psychosis research, such as in studies examining antidepressant response (Bansal et al., 2019) and metabolic patterns across dementia subtypes (L. Su et al., 2016).

In this paper, we leveraged recent advancements in high-resolution 3D-MRSI (A. Klauser et al., 2022), to identify subtle metabolic changes in patients at-risk for psychosis who meet the SCZT or APS criteria, using a whole-brain voxel-based analysis (VBA) approach (Ashburner & Friston, 2000).

The first aim was to demonstrate the reproducibility of the 3D-MRSI technique using independent datasets of healthy controls from 2 different scanning sites by looking at regional metabolic patterns. Second, we aimed to demonstrate the feasibility of performing a whole-brain VBA on 3D-MRSI metabolic maps. Then, we tested the sensitivity of the whole-brain VBA on 3D-MRSI metabolic maps to detect cerebral alterations in a clinical sample of youths at high risk for psychosis, in comparison with more conventional analyses based on structural (sMRI) and diffusion (dMRI) data. Finally, we also tested associations between metabolic levels and subclinical psychotic symptoms or global functioning in patients at-risk for psychosis.

We expected to see a high reproducibility between the two scanning sites as well as metabolic differences between clinical groups and associations with subclinical psychotic symptoms and level of functioning, especially for Glx, tNAA and Ins levels, which were previously shown to be altered in psychosis. Given the modest sample size, we also expected to find no between-group differences and no association with clinical scores using VBA on dMRI or sMRI data.

## 2. Methods

### 2.1 Participants

#### 2.1.1 Lausanne Psychosis Cohort

This study was part of the “Lausanne psychosis cohort” that is currently ongoing in Lausanne, Switzerland (local ethics approval PB_2017-00675). Participants were recruited from the “Treatment and Early Intervention in Psychosis Program” in Lausanne (Baumann et al., 2013), which was expanded with the inclusion of a section for patients at-risk for psychosis. These participants should be between 15 and 35 years old, without any prior psychotic-related diagnosis. Age and sex-matched healthy controls were recruited in parallel to consider the neuroanatomical modifications due to normal brain development in both male and female patients. Controls should be free of any life-time psychiatric diagnosis and should not have any first-relative with a psychosis-related diagnosis. All participants (patients and controls) were assessed at baseline, and then at 6, 12, 24 and 36 months, or at the time of conversion to psychosis.

Patients at risk for psychosis were assessed by the Structured Interview for Psychosis-Risk Syndromes (SIPS (McGlashan et al., 2010; Miller et al., 2002, 2003)). Patients were classified as SCZT if they met a DSM-5 (*Diagnostic and Statistical Manual of Mental Disorders*, 2013) lifetime criteria for schizotypal personality disorder as reported in the SIPS manual. Patients were classified as APS if they met the attenuated positive symptoms with a current progression (i.e. symptoms appeared or worsened during the last year) and with symptoms appearing at least once a week during the last month. All patients and controls underwent demographic and the Global Assessment of Functioning (GAF (American Psychiatric Association, 1998)). Tobacco use was assessed via self-reported daily cigarette consumption, with participants indicating the average number of cigarettes smoked per day. Medication was assessed at each visit and antipsychotics use was reported as chlorpromazine equivalents (Andreasen et al., 2010; Gardner et al., 2010).

Twenty-six patients were included during the time the 3D-MRSI technique was used in the cohort (May 2019 to May 2022). Patients that met both the criteria for APS and SCZT were considered as APS patients since the APS status involves a more recent uprise of symptoms. Therefore, the 26 patients comprised 15 APS and 11 SCZT. All controls with exploitable 3D-MRSI scans were included in the analyses.

#### 2.1.2 Mindfulteen Study (Geneva)

To test the reproducibility of our 3D-MRSI technique in an independent sample scanned at a different location, we recruited the participants from the Mindfulteen study, a randomized controlled crossover clinical trial that assesses the effects of mindfulness-based intervention in healthy adolescents (n=69, 57% females, age =14±0.8). The study included adolescents between 13 and 15 years old, and they were excluded if they had any chronic somatic disease or any significant medical condition, if they underwent psychotherapy in the last 6 months or received psychotropic medication in the past month, and if they met criteria for any psychiatric disorder in the DSM-IV (American Psychiatric Association, 1998) except for anxiety disorder or past major depressive disorder. The study protocol was approved by the Geneva Regional Ethical Committee (CCER 2018–01731). Further details on recruitment, inclusion, and exclusion criteria can be found in the published protocol (Piguet et al., 2022).

### 2.2 Magnetic Resonance scanning

#### 2.2.1 Lausanne

For the Lausanne Psychosis Cohort, magnetic resonance (MR) data were acquired using a 3-Tesla scanner (Magnetom PrismaFit, Siemens Healthineers, Forchheim, Germany) equipped with a 32-channel head coil at the CIBM Center for Biomedical Imaging in Lausanne University Hospital. Each scanning session included first a magnetization-prepared rapid acquisition gradient echo (MPRAGE) T1-weighted sequence, providing a 1 mm in-plane resolution and 1.2 mm slice thickness, with a coverage of 240 × 256 × 160 voxels (i.e. 240 x 256 x 192mm^3^). The sequence parameters were as follows: repetition time (TR) of 2300 ms, echo time (TE) of 2.98 ms, and inversion time (TI) of 900 ms. Time of acquisition was 2 min 58s.

The 3D ¹H-CS-FID-MRSI sequence was described in details by Klauser et al. (A. Klauser et al., 2022). The acquisition had the following parameters: TE of 1.00 ms, TR of 353 ms, flip angle of 40°. The field-of-view (FOV) spanned 210 mm × 160 mm × 105 mm (anterior-posterior, right-left, head-foot directions) with a 95 mm thick slab selection, and a spatial resolution of 5 × 5 × 5.3 mm^3^. The spectral bandwidth was 2 kHz, acquired with a vector size of 512 points. A water reference acquisition was performed with the following parameters: same TE, TR of 25 ms, lower flip angle of 3°, same FOV size, lower resolution of 6.6 x 6.7 x 6.6 mm^3^, same bandwidth and lower vector size of 16 points. Total acquisition time was 22 min, including 2 min for water reference acquisition. The complete Minimum Reporting Standards in MRS (MRSinMRS, (Lin et al., 2021)) are displayed in Table S6.

The diffusion spectrum imaging (DSI) sequence comprised 128 diffusion-weighted images with a maximum b-value of 8000 s/mm², along with one B0 reference image. The acquisition volume was 96 × 96 × 34 voxels with a spatial resolution of 2.2 × 2.2 × 3 mm^3^. The TR and TE for this sequence were 6800 ms and 144 ms, respectively. Acquisition time was 13 min.

#### 2.2.2 Geneva

For the Mindfulteen study, magnetic resonance (MR) data were acquired using a 3-Tesla scanner (Magnetom TrioTim, Siemens Healthineers, Forchheim, Germany) equipped with a 32-channel head coil at the Brain and Behavior Laboratory in Geneva.

The MPRAGE T1-weighted sequence used the same parameters as in the Lausanne psychosis cohort.

Due to hardware differences (i.e., TrioTim vs PrismaFit), the 3D ¹H-CS-FID-MRSI sequence had different parameters than in Lausanne. The acquisition had the following parameters: TE of 1.5 ms, TR of 372 ms and flip angle of 35°. The field-of-view (FOV) was the same as in Lausanne, with the same spatial resolution of 5 × 5 × 5.3 mm^3^, as the spectral bandwidth and vector size. The water reference acquisition had the following parameters: same TE of 1.5 ms, TR of 36 ms, and a lower flip angle of 5°. FOV size, resolution, bandwidth, and vector size were the same as in Lausanne. The complete MRSinMRS are displayed in Table S6.

### 2.3 3D-MRSI reconstruction

Reconstruction was performed using a low-rank model constrained by total generalized variation, with a simultaneous removal of subcutaneous lipid contamination and residual water signals, as described by Klauser et al. (A. Klauser et al., 2024b). Reconstruction benefited from the latest improvements, notably the simultaneous lipid suppression method, in order to address baseline distortion noted in the original publication (A. Klauser et al., 2022). After reconstruction, the spatio-spectral data were analyzed using LCModel (Provencher, 2001) to estimate metabolite levels within each voxel, with the separate water acquisition serving as a reference. LCModel fit setting file is displayed in Table S7. An example of the spectra across the brain of one participant is displayed in Fig. S3.

The LCModel fitting process utilized a basis set comprising the following metabolites: N-acetylaspartate (NAA), N-acetylaspartylglutamate (NAAG), creatine (Cr), phosphocreatine (PCr), glycerophosphocholine (GPC), phosphocholine (PCh), myo-inositol (Ins), scyllo-inositol (sI), glutamate (Glu), glutamine (Gln), lactate (Lac), gamma-aminobutyric acid (GABA), glutathione (GSH), taurine (Tau), aspartate (Asp), and alanine (Ala). Due to the overlap of spectral peaks, only a subset of these metabolites could be reliably resolved. Consequently, some metabolites were combined for analysis: NAA+NAAG into tNAA, Cr+PCr into tCr, GPC+PCh into Cho, Ins, and Glu+Gln into Glx, resulting in five distinct metabolic volumes. The reported metabolite values were neither corrected for tissue-specific T1 weighting nor adjusted for tissue-specific water content. Therefore, they are reported in institutional units (IU) and not in millimolar (mM).

LCModel also generated spectral quality metrics, including the signal-to-noise ratio (SNR), the full-width at half-maximum value of each resolved peak (FWHM) (one value per voxel), and Cramer-Rao Lower Bound (CRLB) (one value per voxel per metabolite).

### 2.4 3D-MRSI selection

#### 2.4.1 Lausanne

Out of the 26 at-risk for psychosis patients scanned in Lausanne, 9 had scans at multiple assessments (2 or 3), totalizing 36 3D-MRSI brain scans. Sixteen controls were scanned, with 5 having scans at multiple assessments, totalizing 23 MRSI brain scans. For each subject metabolic map, we created a quantitative quality mask based on FWHM map, the SNR map and the metabolite CRLB map. These quantitative maps were then combined to generate a binary “quality mask” (Qmask) for each metabolite in each subject. Within each Qmask, voxels were designated reliable (value = 1) if they simultaneously satisfied the following criteria: CRLB of the given metabolite < 20, FWHM < 0.1, and SNR > 4. Voxels failing any of these thresholds were marked as unreliable (value = 0). We finally masked the metabolic maps with these Qmasks. Visual quality check was done by 3 different contributors (EC, PK and AK) to identify presence of motion during acquisition, based on anatomical patterns displayed in the metabolic maps and the global cerebral volume coverage based on the Qmask. From these criteria, we excluded 8 acquisitions from the patients and 5 acquisitions from the controls. This selection brought the number of individual subjects with a usable 3D-MRSI scan to 21 patients (12 APS and 9 SCZT) and 13 controls, totalizing 34 participants for Lausanne. If multiple brain scans were available for a given participant, only the first one was included in the analyses. Spectral quality can be assessed by the mean values of quality metrics in 12 different brain regions (Fig. S2).

Analyses of structural and diffusion MRI data were performed on the same 34 participants.

#### 2.4.2 Geneva

In the Mindfulteen study, the 69 subjects were scanned, 68 of them having 2 or 3 scans at different time points of the study, totalizing 167 3D-MRSI acquisitions. We excluded 53 of them for bad quality, following the same criteria as in the Lausanne psychosis cohort. We then selected the first available brain scan for each given participant. This selection brought the number of individual subjects with a usable 3D-MRSI scan to 61.

### 2.5 3D-MRSI Data Pre-Processing

The preprocessing pipeline normalizes each 3D-MRSI volume to standard space to perform voxel-based analyses. It consists first in a spatial filtering of metabolic spikes, second in a correction of the partial volume effect (PVE), third in a co-registration of the 3D-MRSI map to the subject T1-weighted (T1w) image, and fourth in applying the T1w normalization to the 3D-MRSI co-registered map.

#### 2.5.1 Spatial Filtering

To post-process the metabolic maps derived from 3D-MRSI, we employed an in-house structured denoising and filtering pipeline. A key step involved identifying outlier voxels, referred to as spikes, which are commonly introduced by errors in metabolite quantification using LCModel, particularly when the reference water signal is underestimated. This leads to an overestimation of metabolite levels at those voxels.

Spike detection was performed using a statistical threshold set at the 99th upper end percentile of intensities within the brain mask. No threshold was set for minimum values.

Identified spike voxels were corrected via localized inpainting using a 3×3×3 median filter, ensuring that extreme values were replaced by representative local neighborhood statistics. Additionally, voxels containing NaNs or zero values (interpreted as missing or unreliable estimates within the brain mask) were repaired using biharmonic inpainting (Damelin & Hoang, 2018) followed by localized median filtering to preserve anatomical plausibility. A SpikeMask is generated for each voxel that corresponds to a spike (i.e., voxel value > 99^th^ percentile), which is noted as a 1.

Subsequently, the images were spatially smoothed using a Gaussian kernel with a full width at half maximum (FWHM) of 5 mm, implemented via Nilearn’s smooth_img function (Nilearn contributors et al., 2025). This smoothing scale was chosen to match the intrinsic 5 mm isotropic resolution of the original 3D-MRSI acquisition, ensuring a maximally non-invasive spatial filtering and minimizing the introduction of artificial blurring or interpolation artifacts.

All filtering and inpainting operations were restricted to voxels within the brain mask to avoid contamination from non-brain regions. These spikes were identified and replaced at this stage rather than being later excluded from the analyses using Qmasks. Indeed, their very high values may bias the co-registration with the structural image and contaminate nearby voxels during the interpolation process.

#### 2.5.2 Co-registration with T1w

We used the following pipeline, illustrated in Fig. 1, to normalize each of the 5 metabolic maps to standard space. Briefly, all subject tCr filtered metabolic map were co-registered to each participant’s high-resolution T1w structural skull-striped image (using bet2 from FSL (S. M. Smith, 2002)) using a rigid registration (with mutual information similarity metric) and symmetric diffeomorphic deformation (with cross-correlation similarity metric), as described in Avants et al, with the use of Advanced Normalization Tools, ANTs (Avants et al., 2008).

**Fig. 1:**
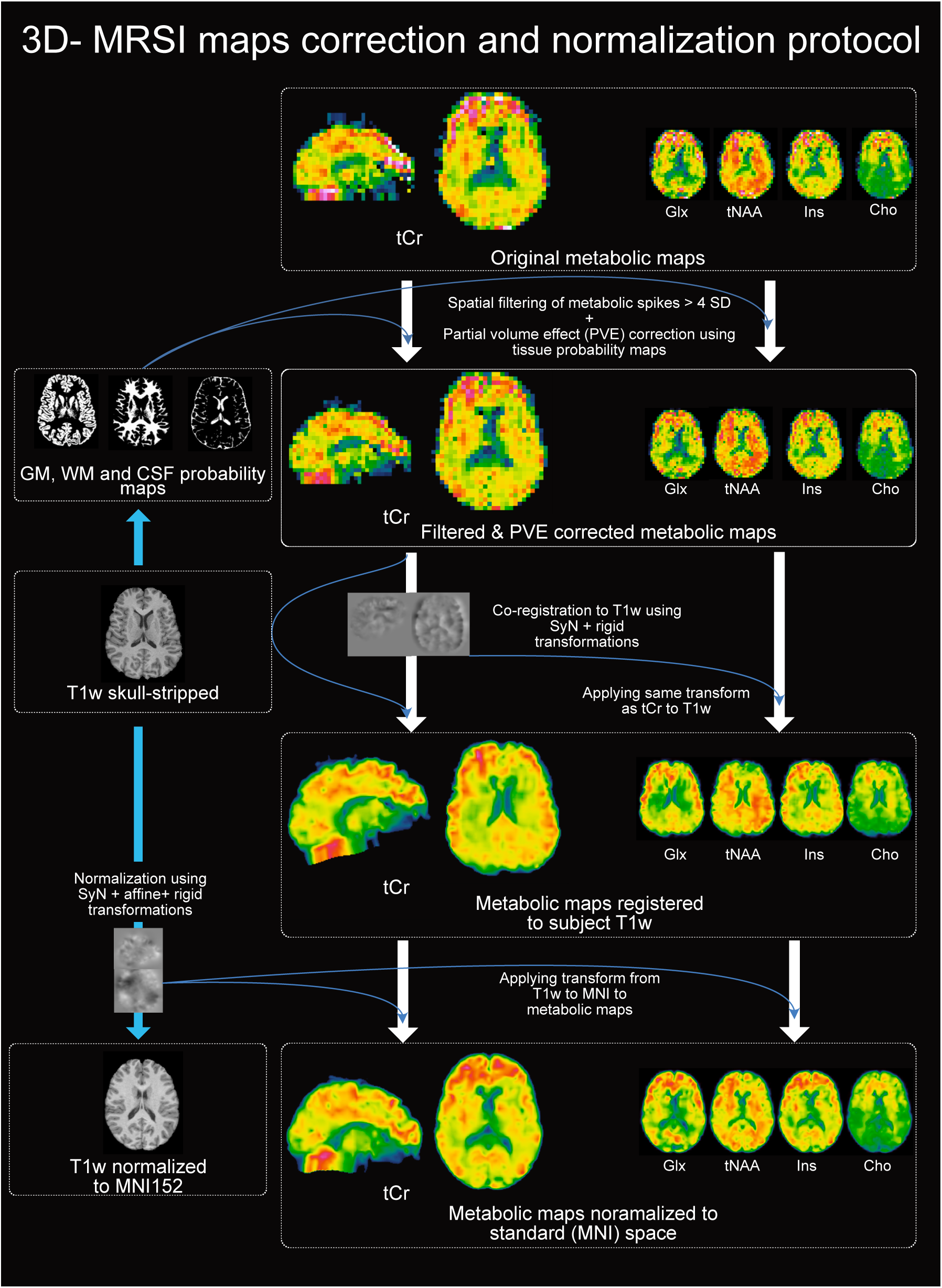
Overview of the correction, co-registration, and normalization pipeline applied to 3D-MRSI data. Original metabolic maps undergo filtering for aberrant values exceeding 4 SD (standard deviation), co-registration to each participant’s T1w anatomical image, PVE (partial volume effect) correction using tissue probability maps down sampled to 3D-MRSI resolution via the inverse co-registration transform, and spatial normalization to MNI152 standard space using the T1w-to-MNI152 transformation. Metabolite distributions may appear slightly different between steps due to volume orientation changes and reslicing. Cho: choline components; Glx: glutamate + glutamine; Ins: Inositol; MNI: Montreal Neurological Institute template; SyN: Symmetric Normalization; T1w: T1-weighted; tCr: total creatine; tNAA: N-acetyl-asparate + N-acetyl-aspartyl-glutamate.

#### 2.5.3 Partial volume effect (PVE) correction

The T1w of each subject was segmented using the Computational Anatomy Toolbox (CAT12, v. 12.9 (Gaser et al., 2024) and Statistical Parametric Mapping (SPM12, v. 7771) on MATLAB (The MathWorks Inc. (2024), v. 24.1.2537033, R2024a). Probability maps of WM, GM and Cerebro-spinal fluid (CSF) were generated for each subject in the T1w space. The inverse transformation from 3D-MRSI to T1w space was applied to these probability maps. Then, we corrected each 3D-MRSI map for GM, WM and CSF partial-volume effect (PVE) using a region-based voxel-wise correction algorithm as implemented in the PETPVC toolbox (Thomas et al., 2016). The 3D-MRSI to T1w transformations were then applied to the 5 PVE corrected 3D-MRSI volumes using tri-linear interpolation.

#### 2.5.4 Normalization

The T1w image of the subject were then spatially normalized to standard space (Montreal Neurological Institue or MNI152 (Fonov et al., 2009)), using a rigid, affine (both with mutual information similarity metric) and deformable symmetric diffeomorphic (with cross-correlation similarity metric) transformations. These transformations were finally applied to the previously co-registered metabolic maps to normalize them to standard space. Each step was visually checked to assure the quality of the registrations and normalizations. Resulting normalized metabolic maps have a spatial resolution of 1 x 1 x 1mm, following guidelines for VBA (Mechelli et al., 2005). Each transformation, from the metabolic 3D-MRSI map space to T1w space to MNI space used a tri-linear interpolation.

#### 2.5.5 Quality masks and metabolite ratio maps

All resolved-metabolite Qmask underwent the same spatial transformations as their corresponding metabolic volumes, using tri-linear interpolation. The SpikeMask of each subject and each metabolite is added to its respective Qmask to exclude the previously identified spikes from the statistical analyses.

To generate metabolite ratio maps, each normalized metabolic map was divided voxel-wise by the sum of all five resolved metabolite level maps. A small constant (ε = 1×10⁻⁶) was added to the denominator to ensure numerical stability. Ratio calculations were performed only within the brain mask.

This entire pipeline is publicly available on Github (https://github.com/MRSI-Psychosis-UP/MRSI-Metabolic-Connectome)

### 2.6 Generalized fractional anisotropy

An automated image correction and processing workflow was applied to individual diffusion-weighted images (DWIs). Specifically, the workflow utilized MRtrix version 3.0.3 and FSL version 6.0.3 to execute several preprocessing steps, including denoising, bias field correction, intensity normalization, head motion correction with corresponding gradient table rotation, as well as eddy current and distortion corrections. Geometric distortions along the phase-encoding direction were addressed using a registration-based correction strategy implemented via Advanced Normalization Tools (ANTs) version 2.4.1. Subsequent to these corrections, Dipy version 1.5.0 was employed to fit second-order diffusion tensors and intravoxel orientation distribution functions (ODFs) through the Simple Harmonic Oscillator-based Reconstruction and Estimation (SHORE) approach (Özarslan et al., 2013). The resulting ODFs, derived from the corrected DWIs, facilitated the calculation of generalized fractional anisotropy (gFA). Analogous to fractional anisotropy (FA), gFA values range from 0 to 1, representing minimal to maximal orientational anisotropy within the ODF. Individual gFA maps were visually inspected to identify and exclude images exhibiting gross abnormalities or artifacts.

### 2.7 Data Analyses

#### 2.7.1 Mean extraction by region

Brain masks in standard space were generated using the Harvard-Oxford atlas (Desikan et al., 2006; Makris et al., 2006): frontal, parietal, temporal and occipital lobes, with subdivisions for grey and white matter as well as thalamus, putamen, pallidum and caudate; each region was also subdivided between left and right side.

A global quality mask was generated for each sample / scanning site (i.e., Geneva and controls from Lausanne), by multiplying all individual Qmasks together. This generated a global Qmask per metabolite per sample. Each regional brain mask (e.g., frontal gray matter) was then multiplied by this global Qmask to extract reliable values for each sample in each brain region. To compare the 2 scanning sites, data from Lausanne were extracted only in controls.

The mean value for each subject was extracted from each of these brain regions. Values were corrected for a different flip-angle across the 2 sites with the following: sin(flip angle (MRSI)) / sin(flip angle (Water acquisition)). To further compare the variation in metabolite levels between the 2 samples, we suppressed any internal reference by computing the metabolite ratio over the sum of the 5 metabolites, as previously described.

Mean values per sample were plotted in R v. 4.4.1 using ggplot2 (Wickham, 2016) and ggridges (Claus O, 2024) packages.

#### 2.7.2 Voxel-based analyses on MRSI data

Voxel-based analyses were performed on normalized 1×1×1mm3 metabolic maps of absolute values, covering both gray and white matter segments, using a generalized linear model (GLM) to test for group differences or variable correlations with the randomise command in FSL software package (v 6.0.7.7). Clusters with corrected P values < .05 (Threshold-Free Cluster Enhancement method (S. Smith & Nichols, 2009) and 10’000 permutations) were considered significant. Each analysis had sex and age as regressor in the GLM.

To address the specific problem of bad quality voxels we used the lesion masking tool implemented in FSL randomise (Winkler et al., 2014): for each subject, their Qmask was used as a voxel-wise regressor, and voxels of poor quality were excluded from the analysis. A grey and white matter mask based on the Harvard-Oxford Atlas was applied to all subjects, to exclude the ventricles and cerebellum from the analyses.

#### 2.7.3 Voxel-based analyses on structural data

T1w images acquired simultaneously with 3D-MRSI data were segmented using the cat12 (Gaser et al., 2024) add-on of SPM12 in 4 ways: grey matter volumes and densities, as well as white matter volumes and densities. A voxel-based analysis using FSL randomise on the whole grey matter or white matter volume were performed on each of these modalities following the analyses done on 3D-MRSI data, using TFCE and 10’000 permutations. Analyses on modulated gray or white matter volumes used total intracranial volume as regressor in the GLM, in addition to sex and age.

#### 2.7.4 Voxel-based analyses on diffusion data

Similarly, whole brain white matter voxel-based analyses on gFA maps were performed using FSL randomise, TFCE and 10’000 permutations. Statistical testing was restricted to white matter by using an inclusive mask (i.e., FMRIB58_FA_1mm template from FSL, thresholded at 2500). Age and sex were used as regressors in the GLM.

#### 2.7.5 Statistical analyses

Statistical analyses were conducted using R version 4.4.2. Group differences in demographics and clinical characteristics, including sex, age, tobacco use in cigarettes per day, GAF scores, and SIPS subscores, were assessed using t-tests for continuous variables and chi-square tests for categorical variables. SIPS scores were summarized as the count of scales within each of the four symptom categories (positive, negative, general, and disorganization) where participants had an intensity score of 3 or higher (on a scale with a maximum score of 6), following guidelines provided by the SIPS manual.

Mean values were extracted from significant clusters identified in voxel-based analyses for each participant and plotted using R (version 4.4.2) with the packages ggplot2 and ggstatsplot (Patil, 2021). For the three-group comparisons (APS vs. SCZT vs. Controls), Holm’s post-hoc tests were applied to the extracted data to correct for multiple comparisons.

## 3. Results

### 3.1 Demographics

A summary of the demographics for the Lausanne participants is displayed in Table 1. Age, sex and tobacco consumption were not significantly different between patients and controls. Global functioning was significantly lower in patients, which was expected. Four patients (2 APS and 2 SCZT) were taking antipsychotic medication at doses ranging from 35 to 240 mg/day of chlorpromazine equivalents, and one of them was also taking escitalopram (20mg/day).

**Table 1:** Demographics and clinical scores for the Lausanne’s sample (young adults at risk for psychosis and controls). APS, Attenuated Psychosis Syndrome; GAF, Global Assessment of Functioning; SCZT, Schizotypal personality disorder; SIPS, Structured Interview for Psychosis-risk Syndromes - number of scales in the category with a score ≥ 3.

| Characteristic | Controls | Patients |  |  | p-value |
| --- | --- | --- | --- | --- | --- |
|  |  | All | APS | SCZT |  |
| Age (Mean±SD) | 23.3 (±3.5) | 22.3 (±4.2) | 23.1 (±4.7) | 21.2 (±3.3) | 0.434 |
| Sex (% females) | 30.8% | 23.8% | 8.3% | 44.4% | 0.713 |
| Tobacco use (% smokers) | 7.7% | 33.3% | 33.3% | 33.3% | 0.119 |
| GAF (Mean±SD) | 85.2 (±2.6) | 47.9 (±9.3) | 46.1 (±9.7) | 50.2 (±8.8) | <0.001 |
| SIPS pos |  | 2.0 (±1.1) | 2.2 (±1.1) | 1.6 (±0.9) |  |
| SIPS neg |  | 2.3 (±1.5) | 2.6 (±1.7) | 2.0 (±1.3) |  |
| SIPS gen |  | 1.2 (±0.9) | 1.0 (±0.7) | 1.4 (±1.1) |  |
| SIPS des |  | 1.2 (±1.0) | 1.5 (±1.1) | 0.8 (±0.8) |  |

### 3.2 Site comparison

Absolute levels of each of the five metabolites (i.e., tCr, Ins, tNAA, Glx, Cho) averaged across all control participants (n = 13 for Lausanne, n = 61 for Geneva) in 12 brain regions, including lobar GM and WM as well as basal ganglia, show a very similar pattern between the 2 scanning sites (Pearson’s r values between 0.94 and 0.96) in Figure 2 (left panel). Normalizing the level of each metabolite (i.e., division by the sum of all 5 metabolites) increases the contrast between brain structures but does not systematically improve the correlations between the 2 scanning sites (Pearson’s r between 0.82 and 0.99) in Figure 2 (right panel). Absolute levels for each brain region are provided in Supplementary Table 1, while normalized levels for the five metabolites are reported in Supplementary Table 2. To facilitate comparison with other studies that typically report metabolite levels normalized to total creatine (tCr), values for Glx, Cho, tNAA, and Ins normalized by tCr levels are presented in Supplementary Table 3 and illustrated in Supplementary Figure 1. There are also strong correlations (Person’s r between 0.9 and 0.98) between the 2 samples for these tCr-normalized values.

**Fig. 2:**
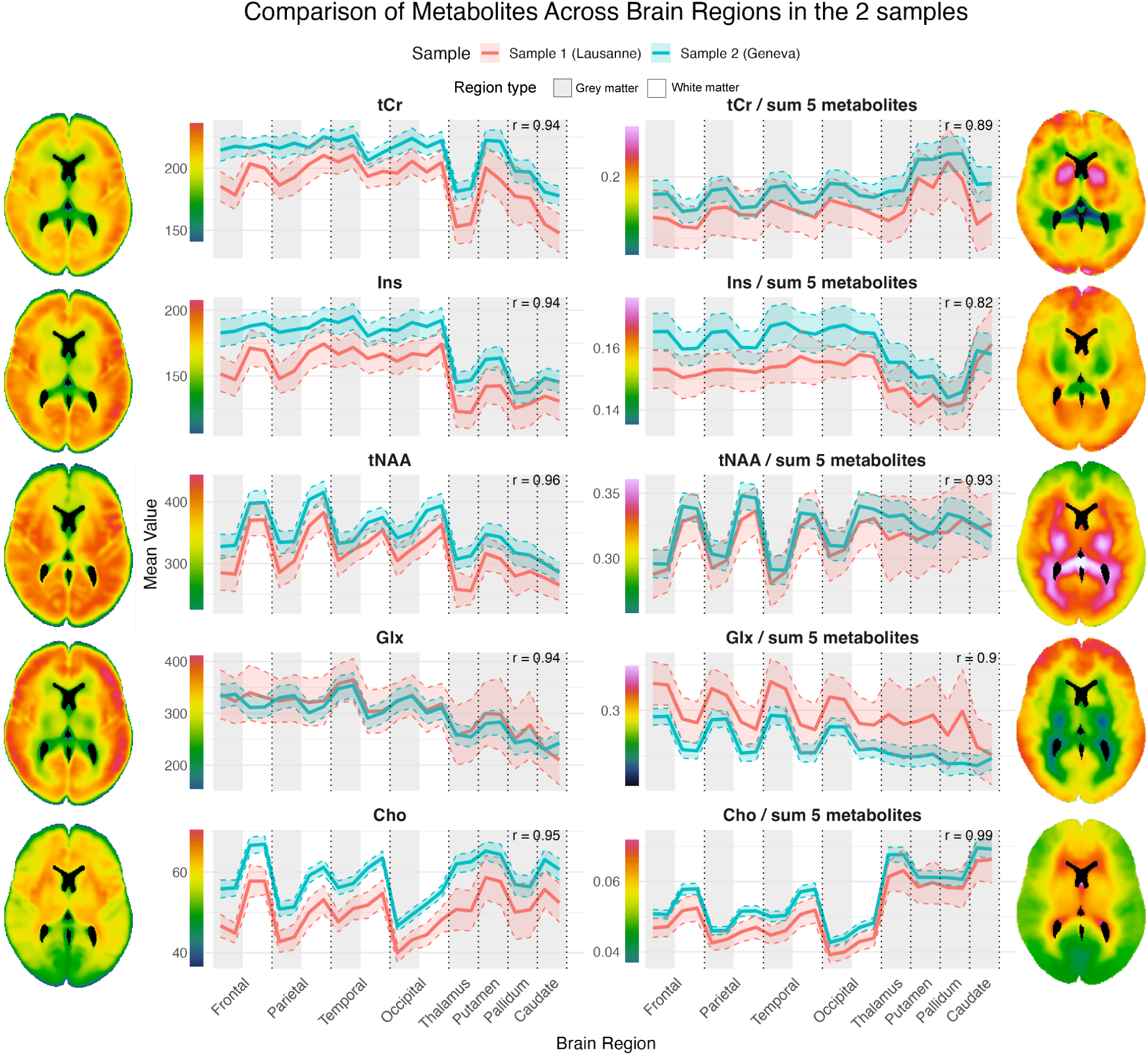
Mean metabolite levels measured across several brain regions (Frontal, Parietal, Temporal, Occipital, divided in grey and white matter; Thalamus, Putamen, Pallidum, Caudate) in participants from two independent samples: Lausanne (controls only, n = 13; red) and Geneva (n = 61; blue). Each region is represented by two data points corresponding to the left and right hemispheres. Absolute metabolite levels (tCr, Ins, tNAA, Glx, Cho; left panel) in IU and respective metabolite ratios relative to the sum of the five metabolites (right panel) are illustrated, accompanied by mean metabolite maps averaged across participants. Correlations between the 2 samples are displayed for each metabolite or metabolite ratio plot on the top right.

### 3.3 Voxel-based analyses

#### 3.3.1 3D-MRSI data

In the Lausanne sample, the comparisons between patients at risk for psychosis (n=21) and controls (n=13) showed a significant cluster of increased tNAA in frontal grey-matter regions especially in the left frontal middle gyrus in patients. (Fig. 3). There was no significant between-group difference for tCr, Ins, Glx or Cho.

**Fig. 3:**
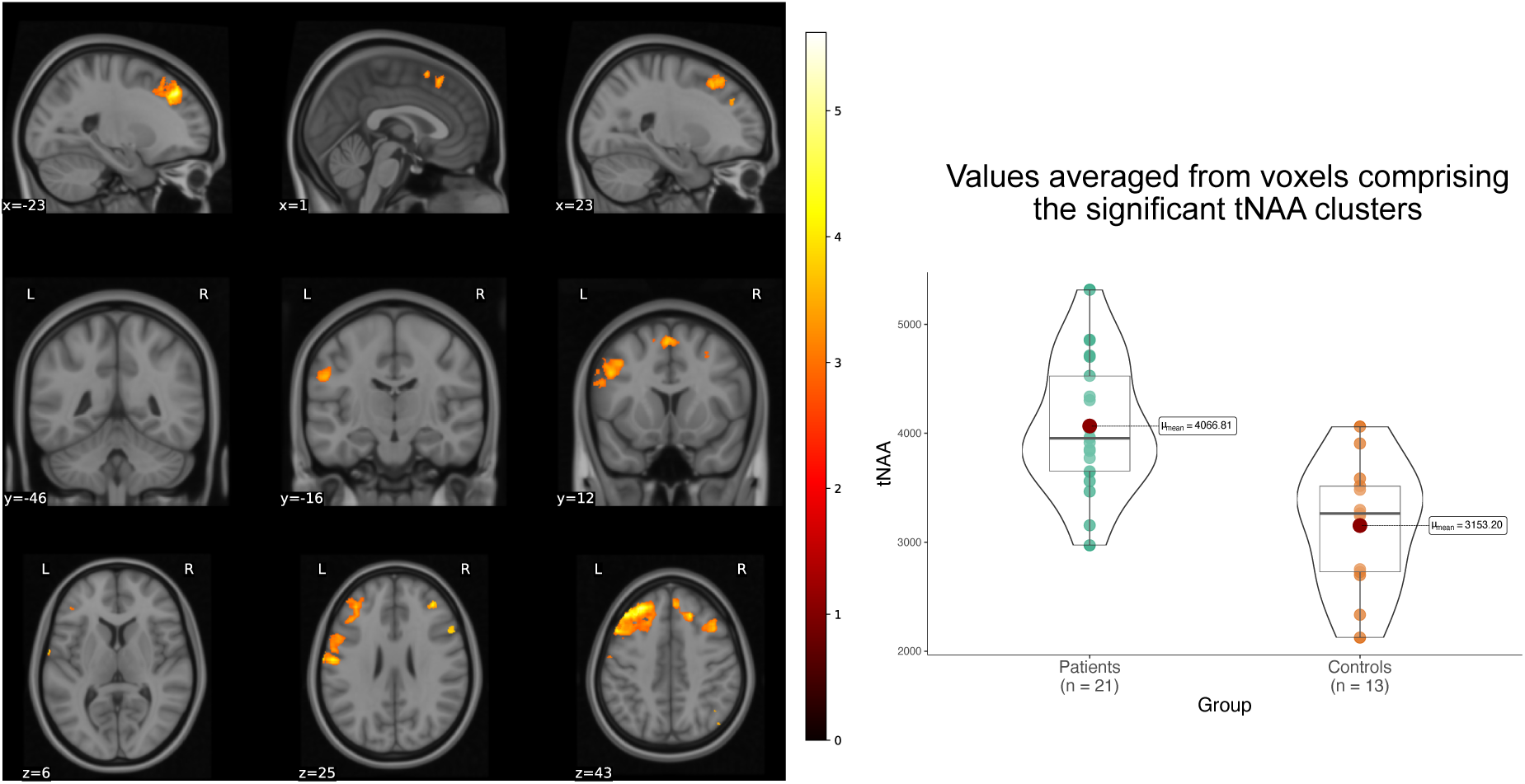
Voxel-based group differences in tNAA between individuals at high risk for psychosis and controls. Increased tNAA levels were observed in frontal grey-matter regions among at-risk individuals, particularly localized within the left middle frontal gyrus. Colored clusters are significant at the whole-brain level (familywise error corrected p < .05). Mean tNAA values extracted from significant clusters are plotted for each group. Analyses are corrected for age and sex. Metabolite levels are reported in IU, and coordinates in MNI152 space.

The 3-group analyses (i.e., SCZT vs APS vs controls) reported significant differences for tNAA, Cho and Ins (Fig. 4). Post-hoc analyses on values averaged from voxels comprising the significant clusters revealed that SCZT patients (n=9) showed the most important increase of tNAA in frontal, parietal and occipital grey-matter compared to controls (n=13). APS patients (n=12) also showed an increased tNAA level compared to controls, but less important than SCZT patients (Fig. 4). Ins was widely increased in grey matter of SCZT patients, including basal ganglia (especially thalamus and caudate) and insula bilaterally. Cho was also increased in SCZT patients in a smaller frontal cluster (Fig. 4). Post-hoc pair-wise comparisons found no increase of Ins or Cho in APS patients when compared to controls.

**Fig. 4:**
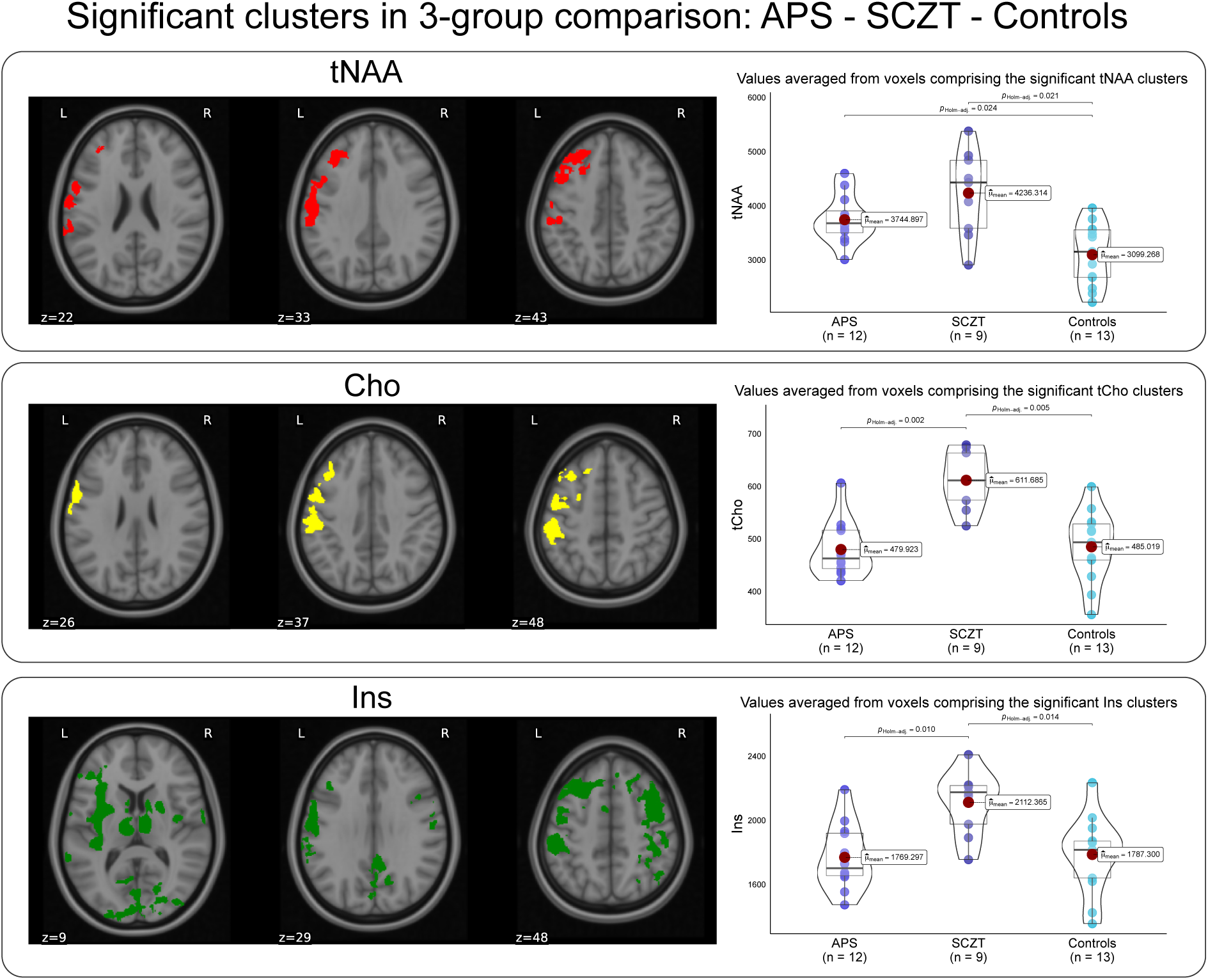
Voxel-based analyses showing differences in tNAA, Ins, and Cho levels among three groups: schizotypal personality disorder (SCZT), attenuated psychosis syndrome (APS), and controls. Compared to controls, increased tNAA levels in SCZT and APS groups were localized primarily to left frontal grey-matter regions with additional small clusters within the left parietal cortex. Cho levels were significantly elevated in SCZT relative to APS and controls, specifically in left frontal and parietal grey matter around the central sulcus. Ins levels were significantly higher in SCZT versus APS and controls, with extensive clusters encompassing frontal, parietal, and occipital grey matter, cingulate cortex, insular regions (predominantly left hemisphere), bilateral thalami, caudate nuclei, and putamina. Colored clusters are significant at the whole-brain level (familywise error corrected p < .05). tNAA, Cho and Ins mean values in these significant clusters are extracted for each participant and plotted. Post-hoc tests using Holm’s test are then performed for between group comparisons. Analyses are corrected for age and sex. Metabolite levels are reported in IU, and coordinates in MNI152 space.

We found no correlation between positive, negative, disorganization and general symptoms from the SIPS and any metabolic measures. Similarly, no correlation was found between GAF and metabolic measures in patients. We also did not find any correlation between tobacco consumption and metabolic levels, in analyses comprising all the participants or only the patients.

#### 3.3.2 Structural data

As expected, we found no significant cluster of gray or white matter volume or density when comparing patients to controls, even when restricting the analyses to brain areas corresponding to the significant tNAA cluster. Similarly, no difference was found in the 3-group analysis between SCZT, APS and controls, and no correlation between positive, negative, disorganization and general symptoms from the SIPS and white or grey matter volume or density.

#### 3.3.3 Diffusion data

Similarly, no cluster of gFA showed significance in the whole brain white matter or in the volume of the significant tNAA cluster for either the 2-group or 3-group analysis, nor for the correlation with the clinical scores.

## 4. Discussion

Improved management of young people suffering from psychosis depends on early detection and intervention, which in turn relies on the prior identification of biological markers of the disorder. The search for brain biomarkers is currently hampered by the limited sensitivity of most brain imaging techniques that are either based solely on the water signal (structural or diffusion MRI) or restricted to a small volume of interest (SVS-MRS). Recent improvements in MRSI now allow high-resolution mapping of multiple metabolites throughout the whole brain volume, and this new approach holds promise for detecting biomarkers in the earliest stages of mental disorders such as in youth at risk for psychosis. In this study we aimed to, first, demonstrate the reproducibility of this 3D-MRSI technique using dataset from 2 different scanning sites; second, demonstrate the feasibility of performing VBA on metabolic maps; and third, test the sensitivity of VBA on metabolic maps on a clinical sample of youths at risk for psychosis, in comparison with more conventional analyses based on structural and diffusion MRI data.

First, mean absolute and relative measures of each of the 5 resolved metabolites across 12 brain regions showed an excellent correlation between the 2 independent samples, despite their intrinsic (i.e., age and sex distribution) and extrinsic (i.e., MRI scanner) differences, demonstrating the replicability of our 3D-MRSI technique. Second, metabolic maps have been successfully pre-processed for VBA (i.e., co-registered with structural images, normalized in standard space, efficiently filtered for signal aberration and corrected for partial volume effect) and analyzed at the voxel and cluster levels, using randomise function in FSL software package. Third, VBA in the Lausanne sample, reported not only an increased left frontal tNAA levels in patients at-risk for psychosis when compared to controls, but also increased tNAA, Ins and Cho levels in a sub-group of patients with SCZT, when compared to APS and controls. As expected, no between-group differences were found using structural (i.e., gray or white matter volumes or densities) or diffusion (i.e., gFA) data, suggesting the higher sensitivity of 3D-MRSI in detecting brain alterations in the early stages of psychosis when compared to structural or diffusion MRI.

### 4.1 Mean measures and previous MRSI literature comparison

To our knowledge, this study is the first to present comprehensive whole-brain 3D-MRSI regional patterns of distributions of metabolites across two independent samples acquired on two different MRI scanners. Regional variations of the metabolites correlate between the 2 samples both on the absolute levels and on the ratio levels. The use of the ratio levels facilitated the comparison of the metabolites between the 2 sites that had different acquisition settings. The cerebellum was excluded from the regional analyses due to incomplete coverage in most participants.

Comparisons with existing literature are primarily possible through relative concentrations. It provides in most cases similar ratios, or similar patterns even in comparison with ultra-high-field MRSI studies. Extensive comparisons were done in the original publication of the 3D-MRSI acquisition and reconstruction protocol (A. Klauser et al., 2022).

Our mean measures align closely with results from Maudsley and colleagues (Maudsley et al., 2009) (acquired using an echo-planar sequence on a 3T Siemens Trio with a TE of 70ms) for tNAA/tCr and Cho/tCr ratios (Ins and Glx were not assessed in this study), and with an antero-posterior gradient for Cho/tCr (Frontal WM: 0.31 in our study vs 0.29, Occipital WM: 0.24 vs 0.22), as well as comparable GM/WM ratios, for example tNAA/tCr (Frontal GM/WM: both 0.85).

Our results also closely match Cho/tCr ratios reported by Goryawala and colleagues (Goryawala et al., 2016) (also acquired using an echo-planar sequence on a 3T Siemens, TE of 17.6 ms), for example: Frontal WM: 0.31 vs 0.26; Parietal WM: 0.27 vs 0.23, Occipital WM: 0.24 vs 0.22. Differences were observed for tNAA/tCr and Glx/tCr ratios. However, the regional patterns remained consistent between studies. Both studies identified higher tNAA/tCr in lobar WM compared to lobar GM (mean GM/WM ratio: 0.86 in our study vs. 0.85) and similarly elevated values in the caudate relative to the thalamus, pallidum, and putamen (e.g., caudate/thalamus ratio: 1.06 vs 1.08).

For Glx/tCr, both studies report higher values in lobar GM compared to lobar WM (GM/WM ratio: 1.15 in our study vs. 1.20), with subcortical regions (caudate, thalamus, pallidum, putamen) showing values similar to lobar WM.

Further, our metabolite ratios align well with recent 7T MRSI studies for example from Hangel and colleagues (Hangel et al., 2021b) (acquired using a free-induction decay rapid concentric ring trajectory sequence on a 7T Siemens Magnetom), notably for Ins/tCr (cortical GM: both 0.86, subcortical WM: 0.87 in our study vs 0.85), and with similar ratios for Cho/tCr, tNAA/Cr and Glx/tCr (notably with the Geneva sample for Glx/tCr). This shows the consistency of our methods with previous literature.

### 4.2 Increase tNAA levels in patients at-risk for psychosis

Our analyses showed increased tNAA levels in the frontal grey matter of patients at-risk for psychosis when compared to controls. The measure of tNAA is the sum of NAA, a marker of neuronal viability previously linked to neuronal loss (Jessen et al., 2001), and NAAG, a neurotransmitter and mGluR3 agonist that inhibits glutamate release and which receptor’s genetic polymorphism has previously been associated with psychosis risk (Harrison et al., 2008; Saini et al., 2017). Most previous SVS-MRS and MRSI studies reported decreased NAA in patients with first episode psychosis or chronic schizophrenia (Paslakis et al., 2014; Yang et al., 2023). On the other hand, meta-analyses on patients at risk for psychosis did not show any significant alteration of tNAA in patients (Romeo et al., 2020; Wang et al., 2020). A specificity of our patient sample is the absence of transition to psychosis during the 3-year follow-up. Our findings of increased tNAA in the frontal regions of patients could therefore be a compensatory mechanism for subjects vulnerable to psychosis who did not experience a first episode psychosis. This may be due to a higher neuronal density in frontal zones or a higher level of NAAG, which regulates excitatory neurotransmission. Increased tNAA levels may also be associated with decreased synaptic pruning, a mechanism that may play a role in psychosis (Glantz & Lewis, 2000; Sekar et al., 2016). Therefore, tNAA could represent a prognosis marker for transition to psychosis, with increased tNAA representing a better outcome. This is also supported by previous studies that found increased tNAA in the frontal cortex as a prognostic marker in patients after their first psychotic episode (Mihaljevic et al., 2024; Wood et al., 2006). However, this hypothesis would need further evaluation and validation in a larger sample that also includes transitions to psychosis.

### 4.3 Increase tNAA and Ins in SCZT subgroup of patients

Our secondary analysis on 2 patients’ subpopulations is based on the differentiation between patients that present a psychopathological state (the APS group) with ones that rather present a pathological trait (the schizotypal personality disorder group). The major difference resides in the stability of the attenuated psychotic symptoms, which are novel or increasing in the last year in the APS group. In this perspective, patients with only the schizotypal trait display stable symptoms for more than one year. Finding the most important tNAA level in this subgroup supports our previous hypothesis of increased tNAA being a compensatory mechanism. In fact, despite having still an important rate of transition to psychosis (i.e., 25-48% (Kirchner et al., 2018)), patients with schizotypal personality disorder have generally a better outcome than patients with SZ and a 60% remission rate (Grilo et al., 2004). Despite the absence of observed differences in volume or gray matter density in these regions, the possibility remains that these results may be associated with the increased cortical thickness observed in the medial orbitofrontal cortex and frontal pole in a large sample of SCZT patients (Kirschner et al., 2022). Altogether, these findings could again point to altered synaptic pruning, reflecting abnormal cortical development.

The increase in Ins levels in widespread grey matter including basal ganglia and insulas in SCZT patients could be an adaptative reaction of glial cells, in which Ins is mostly found, especially in astrocytes (Das et al., 2018; Kim et al., 2005; Rothermundt et al., n.d.), as it is found in brain injuries (Kierans et al., 2014). Previous studies have suggested the important role of astrocytes in brain oxidative regulation and in inflammation (Baxter & Hardingham, 2016; Fernandez-Fernandez et al., 2012), and increased Ins has been consistently found in in brain regions affected by neurodegenerative disorders (Hu et al., 2024; McKiernan et al., 2023; Voevodskaya et al., 2016; Waragai et al., 2017). In that way, increased Ins in patients with chronic symptoms may indicate alterations specific to psychosis. In addition, the functions of regions with increase Ins can be integrated into the psychopathology of psychosis. The insula, involved in affective and somatosensory processing, has been previously reported with reduced volume and dysconnectivity in psychosis (Sheffield et al., 2020; Wylie & Tregellas, 2010).

Ins was increased in striatal structures such as the putamen and caudate, alongside the cingulate gyrus, a closely related structure in the limbic system. These structures have been linked to psychosis due to their important dopaminergic signaling pathways and the mesolimbic action of antipsychotics (McCutcheon et al., 2019).

Finally, the thalamus, especially its anterior and medial parts have been widely discussed in relation to the psychosis spectrum, with an hypoconnectivity to frontal regions (Anticevic & Halassa, 2023; Avram et al., 2021) and a reduced volume, or microstructural alterations in patients suffering from schizophrenia (Alemán-Gómez et al., 2023; Hoang et al., 2021).

Interestingly, the associative striatum, the insula and the thalamus have all been linked to the salience network (H. Huang et al., 2022), which often shows decoupled responses in subjects with psychosis (Wotruba et al., 2014) although this response might be unspecific.

This highlights the possible sensitivity of Ins in previously identified brain structures, although it raises questions about whether these alterations are time- or diagnosis-specific, given that they were only found in SCZT patients.

Beyond its use as a glial marker in brain imaging, Ins is a major astrocytic organic osmolyte that participates in regulatory volume control during sustained osmotic challenges (X. B. Su et al., 2023; Thurston et al., 1989). The widespread elevation of Ins levels that we observed in SCZT may therefore reflect a chronic adaptation of astrocytic osmoregulatory mechanisms in cortical and subcortical regions. However, this remains a mechanistic hypothesis that would require confirmation in studies that directly probe cellular osmotic pathways.

### 4.4 Strengths and limitations

This first study using whole-brain high-resolution 3D-MRSI on patients at-risk for psychosis has several strengths. We developed a novel pipeline that uses state-of-the-art, publicly available tools for maps’ correction, registration and their statistical analyses. This technique provides new insights into the brain metabolism of a highly heterogenous psychiatric population. It offers perspectives not only on what could drive psychosis, but also on what could promote resilience. Its sensitivity in small samples makes it a promising tool for precision psychiatry in specific patient categories. This 3D-MRSI technique has also recently been used to construct a metabolic connectome, in which metabolite similarity gradients and network architecture can be leveraged to unveil brain’s biochemical organization (Lucchetti et al., 2025).

This study has also important limitations. The relatively small size of the sample limits the generalization of the results, and the absence of transition to psychosis in the sample misses an important subcategory of patients that could have been compared to the ones that did not transition. 3D-MRSI results are also difficult to interpret due to the non-specificity of the resolved metabolites and the acquisition of several metabolites that are in the same pathways (N-acetylaspartate and N-acetylaspartyl-glutamate, Glutamine and Glutamate, the different choline components). In addition, metabolite levels are reported in IU, which doesn’t allow their direct comparison between scanning sites and other studies, especially since no phantom calibration and full B1 correction were done. Finally, some MRSI scans had to be ruled out due to global quality issues: this is generally explained by participant movement during a relatively long acquisition (20 minutes). However, this issue will be mitigated in future studies by reducing the acquisition time by about half (A. Klauser et al., 2024b).

Last, it should also be noted that the lipid suppression strategy utilized in our 3D-MRSI technique has the potential to distort the spectrum and could therefore affect the fitting of metabolite signals especially around the 2ppm region, where the tNAA peak is localized (Klauser et al., 2022). According to previous work (Klauser et al., 2024), this distortion is reduced when lipid suppression is integrated into the signal reconstruction, with good reproducibility across measurements and subjects. Consistent with these findings, we also observed good reproducibility between the two samples (Fig. 2). The calculations of coefficients of variation (COV) for tNAA and tCr in patients and controls (Table S9) did not suggest a systematic bias that could solely explain our findings. However, a subtle influence of our lipid suppression method on metabolite measurements, especially on tNAA levels cannot be fully ruled out. Future methodological developments enabling more flexible handling of lipid signals may allow for more direct evaluation of this effect.

## 5. Conclusion

In summary, we were successfully able to use 3D-MRSI and voxel-based analyses to find group differences between subjects at-risk for psychosis (comprising SCZT and APS) and controls, and even differences between SCZT, APS and controls. This opens perspectives for metabolic profiles in this highly heterogenous category of help-seeking patients. We were also able to successfully replicate our preprocessing pipeline in 2 independent samples, paving the way for more extensive studies with 3D-MRSI in clinical and non-clinical populations.

## Ethics

The study protocols were approved by the Geneva and Lausanne Regional Ethical Committees (CCER 2018-01731 and PB_2017-00675 respectively).

## Code availability

The code used to correct and normalize 3D-MRSI maps presented in this paper is available at the following GitHub repository: https://github.com/MRSI-Psychosis-UP/MRSI-Metabolic-Connectome.

## Data availability

Because of the highly sensitive nature of the clinical data used in this study, researchers interested in accessing the dataset should contact the authors with a well-motivated request detailing their research aims and the intended use of the data.

## Authors contribution

Edgar Céléreau: Conceptualization, Methodology, Investigation, Software, Data

Curation, Writing – Original Draft, Review & Editing

Federico Lucchetti: Software, Writing – Review & Editing

Yasser Alemán-Gómez: Formal analysis, Writing – Review & Editing

Daniella Dwir: Supervision, Writing – Review & Editing

Martine Cleusix: Resources, Data Curation, Writing – Review & Editing

Jean-Baptiste Ledoux: Resources, Data Curation, Writing – Review & Editing

Caroline Conchon: Resources, Data Curation, Writing – Review & Editing

Raoul Jenni: Resources, Data Curation, Writing – Review & Editing

Merixtell Bach-Cuadra: Methodology, Writing – Review & Editing

Zoé Schilliger: Resources, Data Curation, Writing – Review & Editing

Alessandra Solida: Resources, Data Curation

Marco Armando: Data Curation, Writing – Review & Editing

Kerstin Jessica Plessen: Funding acquisition, Writing – Review & Editing

Patric Hagmann: Resources, Funding acquisition, Writing – Review & Editing

Philippe Conus: Resources, Funding acquisition, Writing – Review & Editing

Antoine Klauser: Methodology, Software, Validation, Supervision, Writing – Review & Editing

Paul Klauser: Conceptualization, Methodology, Validation, Supervision, Data Curation, Project administration, Funding acquisition, Writing – Review & Editing

## Competing interest

Antoine Klauser is employed by Siemens Healthineers AG, Switzerland. The other authors have nothing to disclose.

## Funding

This study was funded by the Swiss National Science Foundation (grant number 215728). EC was supported by an MD-PhD fellowship from the faculty of biology and medicine, University of Lausanne. PK and DD were sup ported by a fellowship from the Adrian & Simone Frutiger Foundation. Data acquisition in Geneva was supported by a grant from the Leenaards Foundation.

## Acknowledgements

We gratefully acknowledge Camille Piguet, Arnaud Merglen, Frederic Grouiller and all the Mindfulteen team who contributed to the acquisition of the 3D-MRSI dataset in Geneva, along with all participants as well as clinicians from the TIPP program at Lausanne University Hospital.

## Notes

### Competing Interest Statement

AK is employed by Siemens Healthineers AG, Switzerland. The other authors have nothing to disclose.

### Summary of Updates

Modifications made to the introduction. Augmented results. Augmented discussion.

