## Supplementary Material for "High-resolution whole-brain magnetic resonance spectroscopic imaging in youth at risk for psychosis"

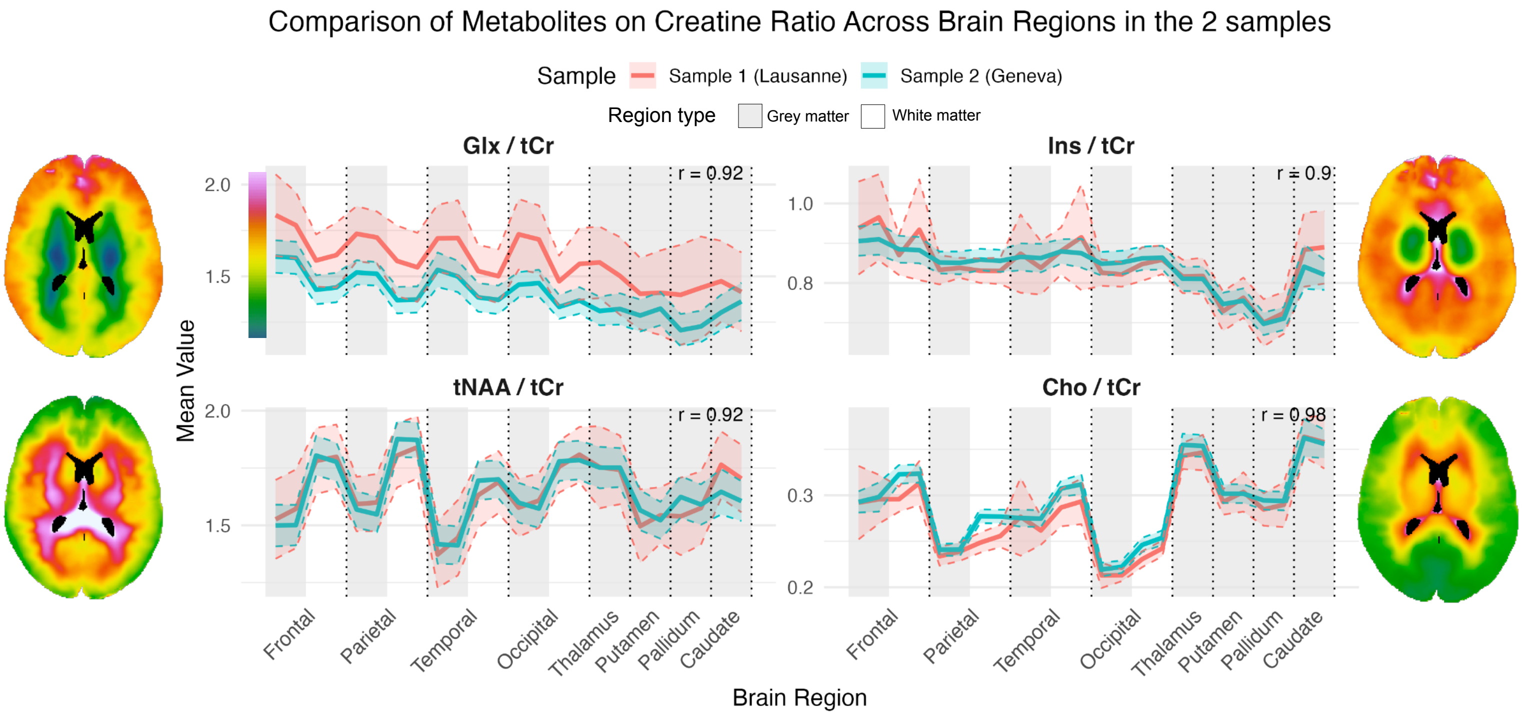


**Fig. S1**: Mean metabolite concentrations normalized on tCr measured across several cerebral regions (Frontal, Parietal, Temporal, Occipital, divided in grey and white matter; Thalamus, Putamen, Pallidum, Caudate) in participants from two independent samples: Lausanne (controls only, n = 13; red) and Geneva (n = 61; blue). Each region is represented by two data points corresponding to the left and right hemispheres. Correlations between the 2 samples are displayed for each metabolite or metabolite ratio plot on the top right.

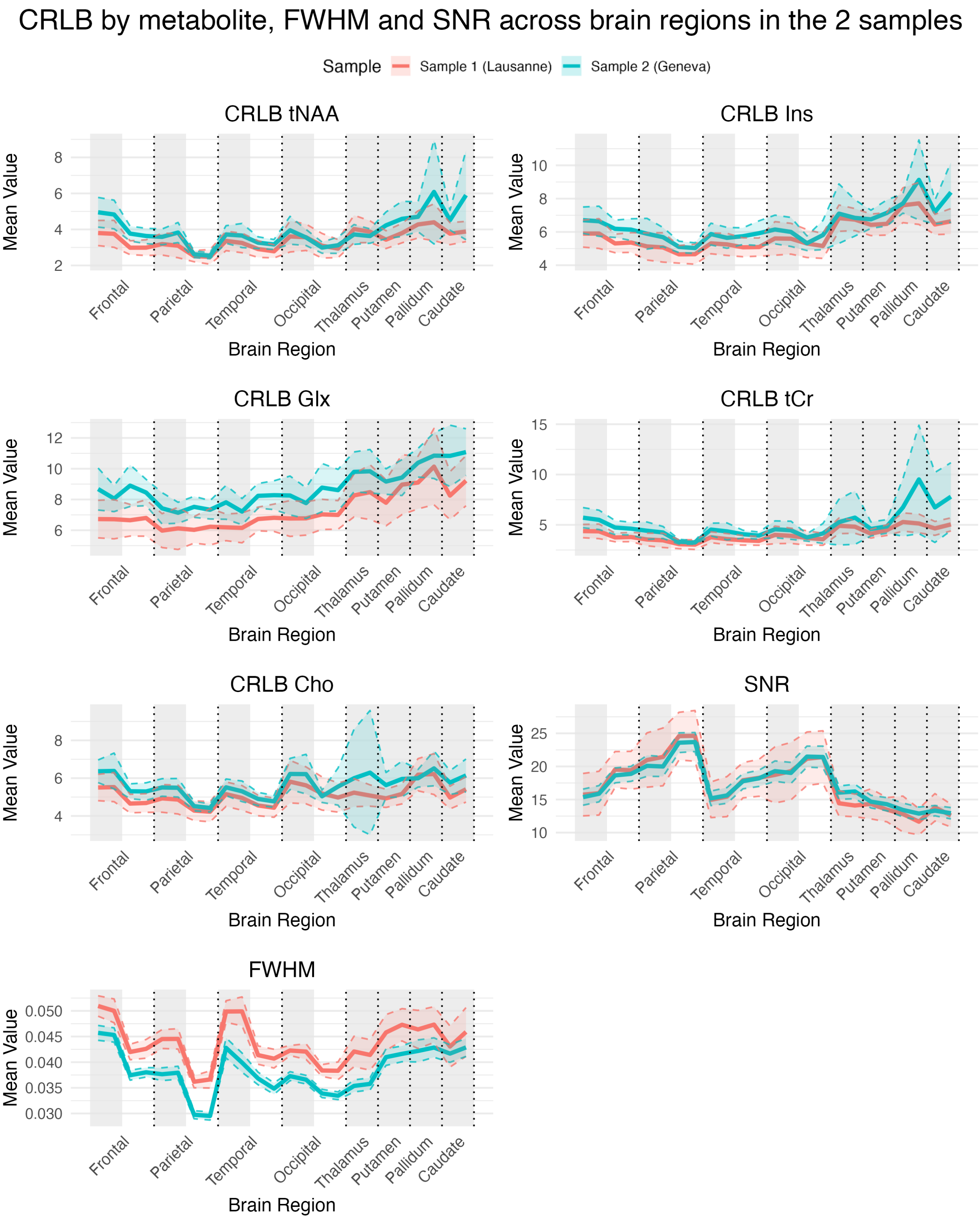


**Fig. S2**: Mean quality indexes (CRLB : Cramer-Rao Lower Bound ; SNR : Signal-to-Noise Ratio ; FWHM : Full Width at Half-Maximum) measured across several cerebral regions (Frontal, Parietal, Temporal, Occipital, divided in grey and white matter ; Thalamus, Putamen, Pallidum, Caudate) in participants from two independent samples : Lausanne (controls only, n = 13; red) and Geneva (n = 61; blue). Each region is represented by two data points corresponding to the left and right hemispheres.

### Spectra example in one subject

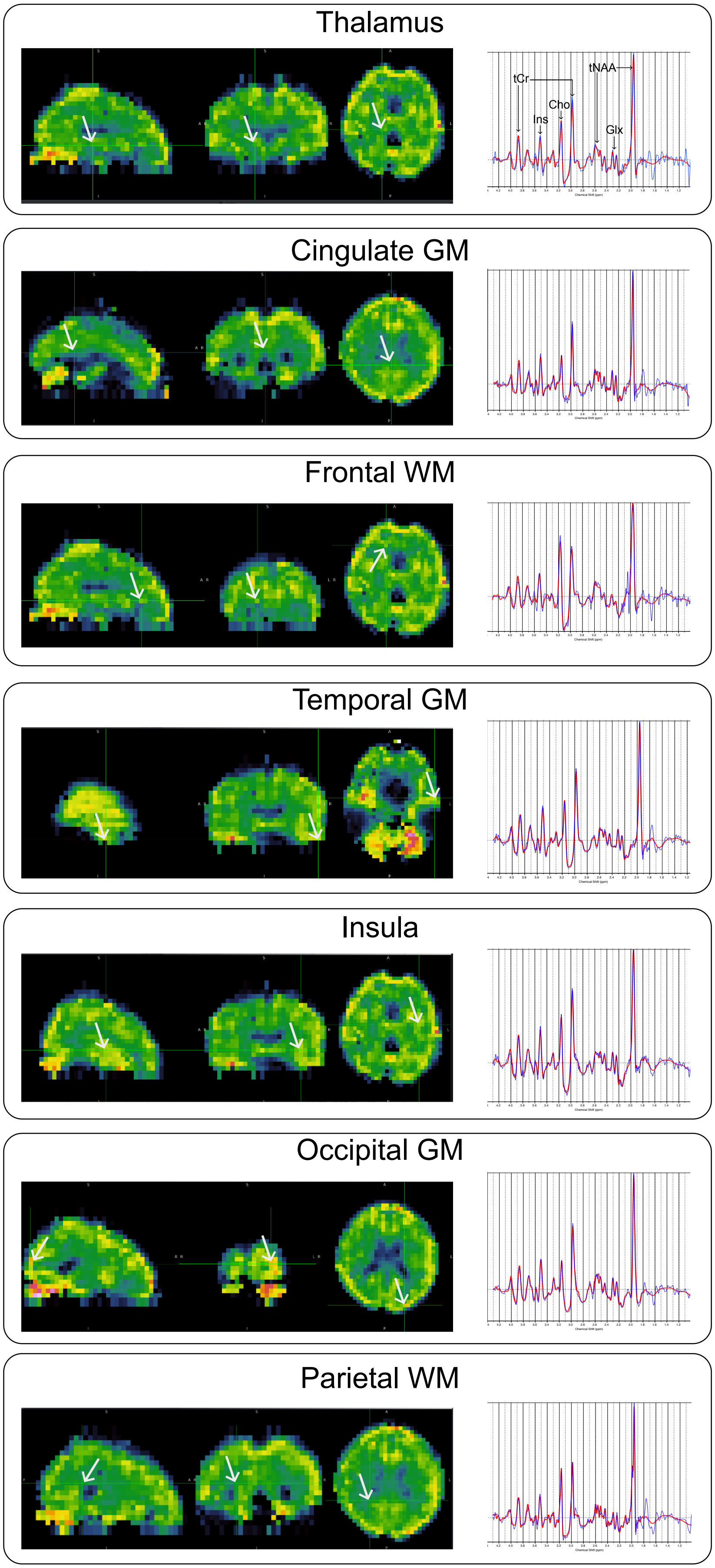


**Fig. S3**: An example of spectra in 7 different voxels of one control subject. Brain region is indicated based on the subject's T1w. Position of the voxel shown on the left, spectra of the given voxel on the right. The blue line corresponds to the reconstructed MRSI spectrum, the red line to the LCModel fit.

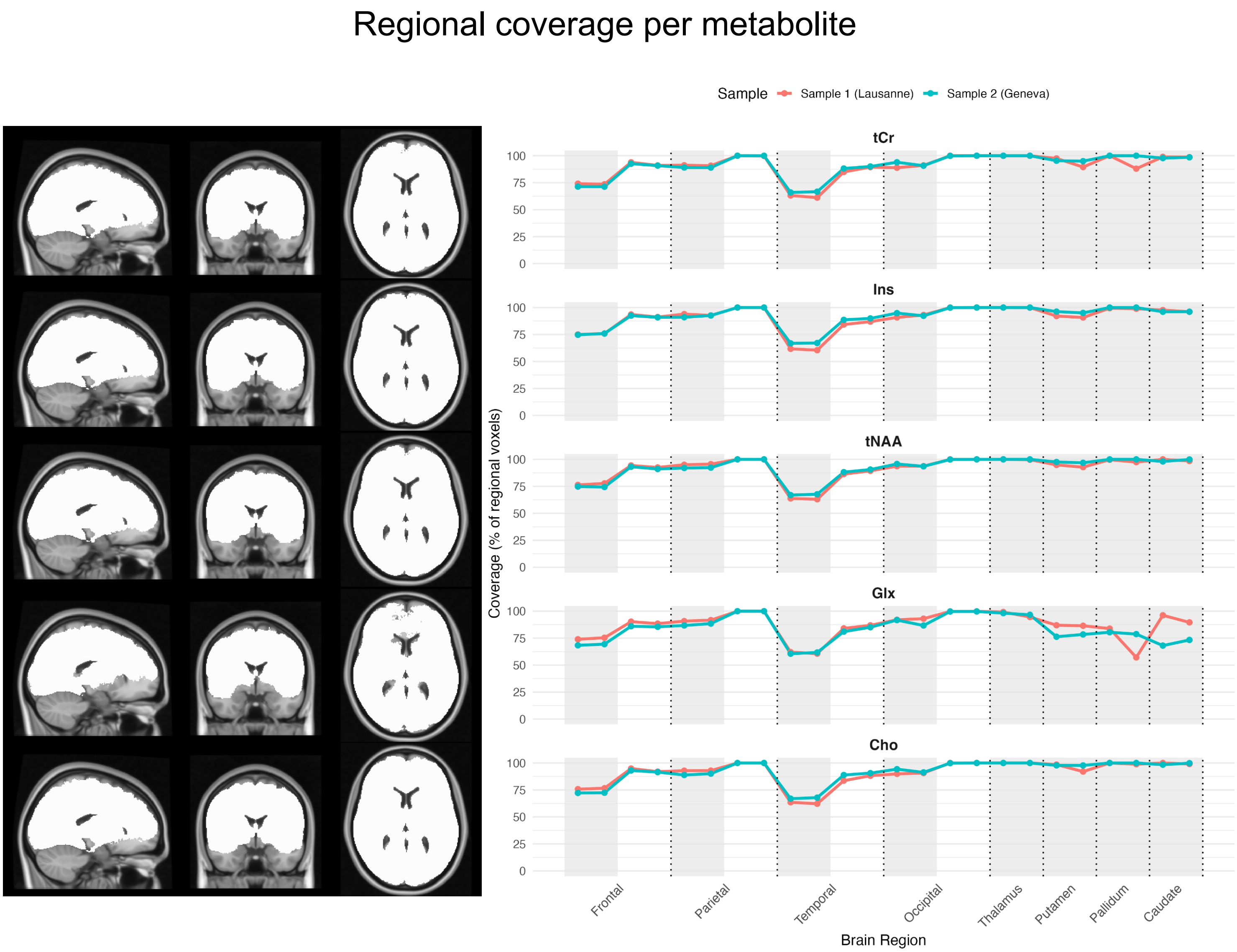


**Fig. S4**: Quality mask population coverage per metabolite. For each metabolite, coverage of the population is displayed in MNI (standard) space on the left. Percentage of each brain region that has good quality is displayed on the left. Frontal GM has a coverage of around 75% and temporal GM has a coverage of around 65%.

#### Table S1. Absolute Metabolic levels by brain regions (corrected for site)

| Brain Region | Type | Side | Cohort | tCr | Cho | Glx | Ins | tNAA |
|---|---|---|---|---|---|---|---|---|
| *Frontal* | *GM* | *L* | *Sample 1 (Lausanne)* | *178.39 ± 19.32* | *44.85 ± 4.07* | *323.45 ± 72.04* | *147.02 ± 20.90* | *282.64 ± 46.95* |
| *Frontal* | *GM* | *L* | *Sample 2 (Geneva)* | *217.64 ± 32.09* | *56.08 ± 7.68* | *336.91 ± 83.91* | *184.07 ± 39.49* | *329.57 ± 71.23* |
| *Frontal* | *GM* | *R* | *Sample 1 (Lausanne)* | *185.41 ± 19.85* | *46.68 ± 4.68* | *336.91 ± 77.59* | *151.48 ± 21.94* | *285.03 ± 46.55* |
| *Frontal* | *GM* | *R* | *Sample 2 (Geneva)* | *214.50 ± 35.12* | *55.82 ± 8.69* | *332.75 ± 89.74* | *182.78 ± 40.77* | *327.56 ± 76.42* |
| *Frontal* | *WM* | *L* | *Sample 1 (Lausanne)* | *200.15 ± 20.00* | *57.76 ± 5.57* | *330.81 ± 82.65* | *169.31 ± 20.96* | *371.07 ± 43.07* |
| *Frontal* | *WM* | *L* | *Sample 2 (Geneva)* | *219.24 ± 27.16* | *66.93 ± 7.69* | *312.17 ± 80.52* | *189.81 ± 36.73* | *399.14 ± 77.01* |
| *Frontal* | *WM* | *R* | *Sample 1 (Lausanne)* | *203.51 ± 22.67* | *57.76 ± 6.73* | *339.88 ± 88.38* | *171.18 ± 23.25* | *370.22 ± 41.74* |
| *Frontal* | *WM* | *R* | *Sample 2 (Geneva)* | *216.39 ± 27.41* | *66.60 ± 8.34* | *311.43 ± 84.02* | *187.97 ± 36.88* | *397.51 ± 77.31* |
| *Parietal* | *GM* | *L* | *Sample 1 (Lausanne)* | *192.43 ± 22.99* | *43.85 ± 5.26* | *328.92 ± 74.74* | *153.77 ± 23.48* | *303.82 ± 49.33* |
| *Parietal* | *GM* | *L* | *Sample 2 (Geneva)* | *220.16 ± 33.58* | *51.35 ± 7.25* | *334.35 ± 84.76* | *185.09 ± 39.35* | *335.56 ± 69.36* |
| *Parietal* | *GM* | *R* | *Sample 1 (Lausanne)* | *186.38 ± 18.14* | *42.71 ± 4.33* | *323.72 ± 67.07* | *147.79 ± 18.75* | *285.47 ± 40.40* |
| *Parietal* | *GM* | *R* | *Sample 2 (Geneva)* | *216.03 ± 33.93* | *50.85 ± 7.27* | *330.00 ± 89.13* | *183.20 ± 38.94* | *334.33 ± 71.77* |
| *Parietal* | *WM* | *L* | *Sample 1 (Lausanne)* | *210.23 ± 23.46* | *53.15 ± 5.80* | *324.11 ± 77.58* | *174.32 ± 24.49* | *382.52 ± 43.20* |

| Brain Region | Type | Side | Cohort | tCr | Cho | Glx | Ins | tNAA |
|---|---|---|---|---|---|---|---|---|
| *Parietal* | *WM* | *L* | *Sample 2 (Geneva)* | *225.08 ± 26.67* | *61.21 ± 6.86* | *312.92 ± 81.94* | *193.26 ± 35.59* | *415.48 ± 71.12* |
| *Parietal* | *WM* | *R* | *Sample 1 (Lausanne)* | *203.11 ± 24.36* | *49.93 ± 6.07* | *320.97 ± 76.01* | *168.51 ± 23.87* | *360.31 ± 39.14* |
| *Parietal* | *WM* | *R* | *Sample 2 (Geneva)* | *216.62 ± 25.60* | *59.07 ± 6.84* | *300.91 ± 81.80* | *186.74 ± 34.16* | *403.38 ± 70.50* |
| *Temporal* | *GM* | *L* | *Sample 1 (Lausanne)* | *210.41 ± 17.05* | *50.93 ± 4.62* | *364.71 ± 67.12* | *171.81 ± 19.34* | *321.97 ± 36.04* |
| *Temporal* | *GM* | *L* | *Sample 2 (Geneva)* | *225.98 ± 30.98* | *57.16 ± 6.87* | *353.88 ± 92.96* | *195.19 ± 38.77* | *336.21 ± 65.39* |
| *Temporal* | *GM* | *R* | *Sample 1 (Lausanne)* | *205.04 ± 15.98* | *47.67 ± 4.15* | *357.06 ± 61.88* | *166.51 ± 17.81* | *305.05 ± 43.56* |
| *Temporal* | *GM* | *R* | *Sample 2 (Geneva)* | *222.09 ± 29.93* | *56.12 ± 6.65* | *347.56 ± 95.05* | *190.68 ± 37.55* | *332.46 ± 68.56* |
| *Temporal* | *WM* | *L* | *Sample 1 (Lausanne)* | *197.42 ± 18.74* | *54.70 ± 5.59* | *304.91 ± 70.34* | *166.75 ± 19.91* | *354.33 ± 26.94* |
| *Temporal* | *WM* | *L* | *Sample 2 (Geneva)* | *212.55 ± 26.32* | *63.52 ± 6.67* | *301.03 ± 82.00* | *185.45 ± 35.56* | *374.65 ± 66.84* |
| *Temporal* | *WM* | *R* | *Sample 1 (Lausanne)* | *193.42 ± 15.68* | *51.81 ± 5.44* | *303.41 ± 68.78* | *163.32 ± 19.53* | *334.44 ± 35.39* |
| *Temporal* | *WM* | *R* | *Sample 2 (Geneva)* | *206.13 ± 23.87* | *61.01 ± 6.22* | *290.95 ± 80.41* | *180.37 ± 33.51* | *366.24 ± 67.91* |
| *Occipital* | *GM* | *L* | *Sample 1 (Lausanne)* | *205.58 ± 16.06* | *43.18 ± 3.78* | *334.88 ± 64.29* | *166.89 ± 16.08* | *323.70 ± 35.12* |
| *Occipital* | *GM* | *L* | *Sample 2 (Geneva)* | *223.97 ± 32.29* | *49.43 ± 5.89* | *333.31 ± 87.29* | *190.68 ± 38.71* | *350.08 ± 73.27* |
| *Occipital* | *GM* | *R* | *Sample 1 (Lausanne)* | *196.15 ± 15.72* | *39.92 ± 3.87* | *322.81 ± 60.49* | *161.15 ± 17.72* | *304.47 ± 42.54* |

| Brain Region | Type | Side | Cohort | tCr | Cho | Glx | Ins | tNAA |
|---|---|---|---|---|---|---|---|---|
| *Occipital* | *GM* | *R* | *Sample 2 (Geneva)* | *218.29 ± 30.45* | *46.63 ± 5.14* | *324.20 ± 87.30* | *184.56 ± 37.09* | *341.57 ± 70.54* |
| *Occipital* | *WM* | *L* | *Sample 1 (Lausanne)* | *204.50 ± 16.73* | *48.16 ± 5.12* | *317.93 ± 65.13* | *174.13 ± 19.78* | *362.84 ± 33.62* |
| *Occipital* | *WM* | *L* | *Sample 2 (Geneva)* | *222.17 ± 28.44* | *55.13 ± 5.86* | *310.49 ± 83.36* | *192.29 ± 37.54* | *393.69 ± 77.14* |
| *Occipital* | *WM* | *R* | *Sample 1 (Lausanne)* | *196.95 ± 14.48* | *44.44 ± 4.84* | *306.07 ± 70.79* | *165.48 ± 20.19* | *341.27 ± 43.49* |
| *Occipital* | *WM* | *R* | *Sample 2 (Geneva)* | *217.06 ± 27.04* | *52.16 ± 5.09* | *300.72 ± 84.01* | *187.58 ± 36.37* | *385.83 ± 73.18* |
| *Thalamus* | *GM* | *L* | *Sample 1 (Lausanne)* | *155.28 ± 25.19* | *50.40 ± 8.23* | *266.94 ± 98.78* | *122.08 ± 20.27* | *255.67 ± 36.75* |
| *Thalamus* | *GM* | *L* | *Sample 2 (Geneva)* | *183.59 ± 28.99* | *62.55 ± 9.04* | *254.40 ± 75.29* | *146.66 ± 30.49* | *312.36 ± 64.70* |
| *Thalamus* | *GM* | *R* | *Sample 1 (Lausanne)* | *152.98 ± 23.08* | *50.74 ± 8.55* | *257.21 ± 94.62* | *123.04 ± 19.80* | *257.98 ± 48.22* |
| *Thalamus* | *GM* | *R* | *Sample 2 (Geneva)* | *181.19 ± 27.59* | *62.02 ± 8.92* | *259.45 ± 75.49* | *145.00 ± 29.39* | *307.10 ± 62.87* |
| *Putamen* | *GM* | *L* | *Sample 1 (Lausanne)* | *190.52 ± 27.98* | *57.67 ± 9.63* | *298.10 ± 113.82* | *142.50 ± 24.69* | *307.61 ± 39.69* |
| *Putamen* | *GM* | *L* | *Sample 2 (Geneva)* | *221.59 ± 36.01* | *64.36 ± 10.17* | *283.20 ± 94.01* | *163.60 ± 35.96* | *343.00 ± 75.13* |
| *Putamen* | *GM* | *R* | *Sample 1 (Lausanne)* | *200.21 ± 30.08* | *58.63 ± 9.16* | *299.70 ± 102.73* | *142.01 ± 21.42* | *316.07 ± 46.84* |
| *Putamen* | *GM* | *R* | *Sample 2 (Geneva)* | *222.41 ± 31.51* | *65.23 ± 9.02* | *280.46 ± 91.67* | *162.54 ± 31.05* | *347.28 ± 71.69* |
| *Pallidum* | *GM* | *L* | *Sample 1 (Lausanne)* | *175.92 ± 31.99* | *50.66 ± 11.48* | *277.16 ± 106.45* | *128.79 ± 28.20* | *287.19 ± 45.43* |

| Brain Region | Type | Side | Cohort | tCr | Cho | Glx | Ins | tNAA |
|---|---|---|---|---|---|---|---|---|
| *Pallidum* | *GM* | *L* | *Sample 2 (Geneva)* | *196.95 ± 37.52* | *56.33 ± 10.51* | *248.62 ± 85.23* | *137.69 ± 32.37* | *313.46 ± 81.46* |
| *Pallidum* | *GM* | *R* | *Sample 1 (Lausanne)* | *177.46 ± 39.89* | *50.05 ± 11.20* | *250.81 ± 95.30* | *125.39 ± 27.27* | *279.54 ± 54.65* |
| *Pallidum* | *GM* | *R* | *Sample 2 (Geneva)* | *197.62 ± 34.96* | *56.92 ± 10.27* | *243.32 ± 92.00* | *136.83 ± 30.95* | *317.13 ± 80.47* |
| *Caudate* | *GM* | *L* | *Sample 1 (Lausanne)* | *147.73 ± 25.39* | *52.38 ± 8.00* | *210.12 ± 79.21* | *130.68 ± 24.08* | *264.74 ± 40.58* |
| *Caudate* | *GM* | *L* | *Sample 2 (Geneva)* | *177.73 ± 30.25* | *60.65 ± 10.46* | *242.94 ± 84.53* | *145.28 ± 32.70* | *285.40 ± 65.49* |
| *Caudate* | *GM* | *R* | *Sample 1 (Lausanne)* | *154.98 ± 25.16* | *55.70 ± 7.50* | *230.46 ± 73.17* | *134.53 ± 20.64* | *276.67 ± 46.50* |
| *Caudate* | *GM* | *R* | *Sample 2 (Geneva)* | *180.80 ± 30.02* | *63.09 ± 9.84* | *230.17 ± 66.48* | *148.25 ± 30.41* | *299.67 ± 63.27* |

#### Table S2. Ratio of metabolite / sum of 5 metabolites by brain region

| Brain Region | Type | Side | Cohort | tCr / sum | Cho / sum | Glx / sum | Ins / sum | tNAA / sum |
|---|---|---|---|---|---|---|---|---|
| *Frontal* | *GM* | *L* | *Sample 1 (Lausanne)* | *0.18 ± 0.02* | *0.05 ± 0.00* | *0.33 ± 0.04* | *0.15 ± 0.01* | *0.29 ± 0.03* |
| *Frontal* | *GM* | *L* | *Sample 2 (Geneva)* | *0.19 ± 0.02* | *0.05 ± 0.00* | *0.29 ± 0.04* | *0.17 ± 0.02* | *0.30 ± 0.04* |
| *Frontal* | *GM* | *R* | *Sample 1 (Lausanne)* | *0.18 ± 0.02* | *0.05 ± 0.00* | *0.33 ± 0.04* | *0.15 ± 0.01* | *0.29 ± 0.03* |
| *Frontal* | *GM* | *R* | *Sample 2 (Geneva)* | *0.19 ± 0.02* | *0.05 ± 0.00* | *0.29 ± 0.04* | *0.17 ± 0.02* | *0.30 ± 0.04* |
| *Frontal* | *WM* | *L* | *Sample 1 (Lausanne)* | *0.18 ± 0.01* | *0.05 ± 0.01* | *0.29 ± 0.04* | *0.15 ± 0.01* | *0.33 ± 0.03* |
| *Frontal* | *WM* | *L* | *Sample 2 (Geneva)* | *0.19 ± 0.02* | *0.06 ± 0.01* | *0.26 ± 0.03* | *0.16 ± 0.02* | *0.34 ± 0.04* |
| *Frontal* | *WM* | *R* | *Sample 1 (Lausanne)* | *0.18 ± 0.01* | *0.05 ± 0.01* | *0.29 ± 0.04* | *0.15 ± 0.01* | *0.33 ± 0.03* |
| *Frontal* | *WM* | *R* | *Sample 2 (Geneva)* | *0.19 ± 0.02* | *0.06 ± 0.01* | *0.26 ± 0.04* | *0.16 ± 0.02* | *0.34 ± 0.04* |
| *Parietal* | *GM* | *L* | *Sample 1 (Lausanne)* | *0.19 ± 0.02* | *0.04 ± 0.00* | *0.32 ± 0.04* | *0.15 ± 0.01* | *0.30 ± 0.03* |
| *Parietal* | *GM* | *L* | *Sample 2 (Geneva)* | *0.20 ± 0.02* | *0.05 ± 0.00* | *0.29 ± 0.04* | *0.17 ± 0.02* | *0.30 ± 0.04* |
| *Parietal* | *GM* | *R* | *Sample 1 (Lausanne)* | *0.19 ± 0.02* | *0.04 ± 0.00* | *0.32 ± 0.04* | *0.15 ± 0.01* | *0.29 ± 0.03* |
| *Parietal* | *GM* | *R* | *Sample 2 (Geneva)* | *0.19 ± 0.02* | *0.05 ± 0.00* | *0.29 ± 0.04* | *0.17 ± 0.02* | *0.30 ± 0.04* |
| *Parietal* | *WM* | *L* | *Sample 1 (Lausanne)* | *0.18 ± 0.02* | *0.05 ± 0.01* | *0.28 ± 0.04* | *0.15 ± 0.01* | *0.34 ± 0.03* |

| Brain Region | Type | Side | Cohort | tCr / sum | Cho / sum | Glx / sum | Ins / sum | tNAA / sum |
|---|---|---|---|---|---|---|---|---|
| *Parietal* | *WM* | *L* | *Sample 2 (Geneva)* | *0.19 ± 0.02* | *0.05 ± 0.01* | *0.25 ± 0.04* | *0.16 ± 0.02* | *0.35 ± 0.04* |
| *Parietal* | *WM* | *R* | *Sample 1 (Lausanne)* | *0.18 ± 0.02* | *0.05 ± 0.00* | *0.29 ± 0.04* | *0.15 ± 0.01* | *0.33 ± 0.03* |
| *Parietal* | *WM* | *R* | *Sample 2 (Geneva)* | *0.19 ± 0.02* | *0.05 ± 0.01* | *0.25 ± 0.04* | *0.16 ± 0.02* | *0.35 ± 0.04* |
| *Temporal* | *GM* | *L* | *Sample 1 (Lausanne)* | *0.19 ± 0.02* | *0.05 ± 0.00* | *0.32 ± 0.04* | *0.15 ± 0.01* | *0.29 ± 0.03* |
| *Temporal* | *GM* | *L* | *Sample 2 (Geneva)* | *0.20 ± 0.02* | *0.05 ± 0.01* | *0.29 ± 0.04* | *0.17 ± 0.02* | *0.29 ± 0.04* |
| *Temporal* | *GM* | *R* | *Sample 1 (Lausanne)* | *0.19 ± 0.02* | *0.04 ± 0.00* | *0.33 ± 0.04* | *0.15 ± 0.01* | *0.28 ± 0.03* |
| *Temporal* | *GM* | *R* | *Sample 2 (Geneva)* | *0.20 ± 0.02* | *0.05 ± 0.01* | *0.29 ± 0.04* | *0.17 ± 0.02* | *0.29 ± 0.04* |
| *Temporal* | *WM* | *L* | *Sample 1 (Lausanne)* | *0.18 ± 0.01* | *0.05 ± 0.01* | *0.28 ± 0.05* | *0.16 ± 0.01* | *0.33 ± 0.03* |
| *Temporal* | *WM* | *L* | *Sample 2 (Geneva)* | *0.19 ± 0.02* | *0.06 ± 0.01* | *0.25 ± 0.04* | *0.16 ± 0.02* | *0.33 ± 0.04* |
| *Temporal* | *WM* | *R* | *Sample 1 (Lausanne)* | *0.19 ± 0.01* | *0.05 ± 0.01* | *0.28 ± 0.05* | *0.16 ± 0.01* | *0.32 ± 0.04* |
| *Temporal* | *WM* | *R* | *Sample 2 (Geneva)* | *0.19 ± 0.02* | *0.06 ± 0.01* | *0.25 ± 0.04* | *0.17 ± 0.02* | *0.33 ± 0.04* |
| *Occipital* | *GM* | *L* | *Sample 1 (Lausanne)* | *0.19 ± 0.01* | *0.04 ± 0.00* | *0.31 ± 0.04* | *0.15 ± 0.01* | *0.31 ± 0.03* |
| *Occipital* | *GM* | *L* | *Sample 2 (Geneva)* | *0.20 ± 0.02* | *0.04 ± 0.00* | *0.28 ± 0.04* | *0.17 ± 0.02* | *0.31 ± 0.04* |
| *Occipital* | *GM* | *R* | *Sample 1 (Lausanne)* | *0.19 ± 0.01* | *0.04 ± 0.00* | *0.31 ± 0.04* | *0.16 ± 0.01* | *0.30 ± 0.03* |

| **Brain Region** | **Type** | **Side** | **Cohort** | **tCr / sum** | **Cho / sum** | **Glx / sum** | **Ins / sum** | **tNAA / sum** |
|---|---|---|---|---|---|---|---|---|
| *Occipital* | *GM* | *R* | *Sample 2 (Geneva)* | *0.20 ± 0.02* | *0.04 ± 0.00* | *0.28 ± 0.04* | *0.17 ± 0.02* | *0.31 ± 0.04* |
| *Occipital* | *WM* | *L* | *Sample 1 (Lausanne)* | *0.18 ± 0.01* | *0.04 ± 0.00* | *0.28 ± 0.04* | *0.16 ± 0.01* | *0.33 ± 0.03* |
| *Occipital* | *WM* | *L* | *Sample 2 (Geneva)* | *0.19 ± 0.02* | *0.05 ± 0.01* | *0.26 ± 0.04* | *0.16 ± 0.02* | *0.34 ± 0.04* |
| *Occipital* | *WM* | *R* | *Sample 1 (Lausanne)* | *0.19 ± 0.01* | *0.04 ± 0.00* | *0.29 ± 0.05* | *0.16 ± 0.01* | *0.33 ± 0.04* |
| *Occipital* | *WM* | *R* | *Sample 2 (Geneva)* | *0.19 ± 0.02* | *0.05 ± 0.01* | *0.26 ± 0.04* | *0.17 ± 0.02* | *0.34 ± 0.05* |
| *Thalamus* | *GM* | *L* | *Sample 1 (Lausanne)* | *0.19 ± 0.02* | *0.06 ± 0.01* | *0.28 ± 0.06* | *0.15 ± 0.01* | *0.32 ± 0.05* |
| *Thalamus* | *GM* | *L* | *Sample 2 (Geneva)* | *0.19 ± 0.02* | *0.07 ± 0.01* | *0.25 ± 0.04* | *0.16 ± 0.02* | *0.33 ± 0.04* |
| *Thalamus* | *GM* | *R* | *Sample 1 (Lausanne)* | *0.18 ± 0.01* | *0.06 ± 0.01* | *0.30 ± 0.07* | *0.15 ± 0.01* | *0.31 ± 0.05* |
| *Thalamus* | *GM* | *R* | *Sample 2 (Geneva)* | *0.19 ± 0.02* | *0.07 ± 0.01* | *0.25 ± 0.04* | *0.16 ± 0.02* | *0.33 ± 0.05* |
| *Putamen* | *GM* | *L* | *Sample 1 (Lausanne)* | *0.20 ± 0.02* | *0.06 ± 0.01* | *0.29 ± 0.07* | *0.14 ± 0.01* | *0.32 ± 0.04* |
| *Putamen* | *GM* | *L* | *Sample 2 (Geneva)* | *0.21 ± 0.02* | *0.06 ± 0.01* | *0.25 ± 0.04* | *0.15 ± 0.02* | *0.32 ± 0.04* |
| *Putamen* | *GM* | *R* | *Sample 1 (Lausanne)* | *0.20 ± 0.02* | *0.06 ± 0.01* | *0.29 ± 0.06* | *0.14 ± 0.01* | *0.31 ± 0.05* |
| *Putamen* | *GM* | *R* | *Sample 2 (Geneva)* | *0.21 ± 0.03* | *0.06 ± 0.01* | *0.25 ± 0.05* | *0.15 ± 0.02* | *0.32 ± 0.05* |
| *Pallidum* | *GM* | *L* | *Sample 1 (Lausanne)* | *0.20 ± 0.02* | *0.06 ± 0.01* | *0.30 ± 0.08* | *0.14 ± 0.01* | *0.33 ± 0.05* |

| Brain Region | Type | Side | Cohort | tCr / sum | Cho / sum | Glx / sum | Ins / sum | tNAA / sum |
|---|---|---|---|---|---|---|---|---|
| *Pallidum* | *GM* | *L* | *Sample 2 (Geneva)* | *0.21 ± 0.03* | *0.06 ± 0.01* | *0.24 ± 0.04* | *0.15 ± 0.02* | *0.33 ± 0.05* |
| *Pallidum* | *GM* | *R* | *Sample 1 (Lausanne)* | *0.21 ± 0.02* | *0.06 ± 0.01* | *0.27 ± 0.06* | *0.14 ± 0.01* | *0.32 ± 0.05* |
| *Pallidum* | *GM* | *R* | *Sample 2 (Geneva)* | *0.21 ± 0.03* | *0.06 ± 0.01* | *0.24 ± 0.05* | *0.14 ± 0.02* | *0.33 ± 0.05* |
| *Caudate* | *GM* | *L* | *Sample 1 (Lausanne)* | *0.19 ± 0.02* | *0.07 ± 0.01* | *0.25 ± 0.05* | *0.16 ± 0.02* | *0.33 ± 0.04* |
| *Caudate* | *GM* | *L* | *Sample 2 (Geneva)* | *0.20 ± 0.03* | *0.07 ± 0.01* | *0.25 ± 0.05* | *0.16 ± 0.03* | *0.32 ± 0.05* |
| *Caudate* | *GM* | *R* | *Sample 1 (Lausanne)* | *0.18 ± 0.02* | *0.07 ± 0.01* | *0.26 ± 0.05* | *0.16 ± 0.02* | *0.32 ± 0.04* |
| *Caudate* | *GM* | *R* | *Sample 2 (Geneva)* | *0.20 ± 0.03* | *0.07 ± 0.01* | *0.24 ± 0.04* | *0.16 ± 0.02* | *0.32 ± 0.05* |

#### Table S3. Ratio of metabolite / creatine by brain region

| Brain region | Type | Side | Cohort | Cho / tCr | Glx / tCr | Ins / tCr | tNAA / tCr |
|---|---|---|---|---|---|---|---|
| *Frontal* | *GM* | *L* | *Sample 1 (Lausanne)* | *0.30 ± 0.04* | *1.78 ± 0.25* | *0.96 ± 0.17* | *1.57 ± 0.27* |
| *Frontal* | *GM* | *L* | *Sample 2 (Geneva)* | *0.30 ± 0.06* | *1.60 ± 0.33* | *0.91 ± 0.15* | *1.50 ± 0.34* |
| *Frontal* | *GM* | *R* | *Sample 1 (Lausanne)* | *0.29 ± 0.06* | *1.83 ± 0.33* | *0.94 ± 0.18* | *1.53 ± 0.27* |
| *Frontal* | *GM* | *R* | *Sample 2 (Geneva)* | *0.29 ± 0.04* | *1.61 ± 0.34* | *0.91 ± 0.14* | *1.50 ± 0.35* |
| *Frontal* | *WM* | *L* | *Sample 1 (Lausanne)* | *0.31 ± 0.04* | *1.62 ± 0.28* | *0.93 ± 0.21* | *1.80 ± 0.23* |
| *Frontal* | *WM* | *L* | *Sample 2 (Geneva)* | *0.32 ± 0.04* | *1.44 ± 0.31* | *0.88 ± 0.13* | *1.78 ± 0.33* |
| *Frontal* | *WM* | *R* | *Sample 1 (Lausanne)* | *0.30 ± 0.02* | *1.59 ± 0.22* | *0.87 ± 0.08* | *1.78 ± 0.24* |
| *Frontal* | *WM* | *R* | *Sample 2 (Geneva)* | *0.32 ± 0.04* | *1.43 ± 0.30* | *0.88 ± 0.13* | *1.80 ± 0.34* |
| *Parietal* | *GM* | *L* | *Sample 1 (Lausanne)* | *0.24 ± 0.02* | *1.71 ± 0.22* | *0.84 ± 0.04* | *1.60 ± 0.21* |
| *Parietal* | *GM* | *L* | *Sample 2 (Geneva)* | *0.24 ± 0.03* | *1.51 ± 0.25* | *0.85 ± 0.11* | *1.55 ± 0.30* |
| *Parietal* | *GM* | *R* | *Sample 1 (Lausanne)* | *0.23 ± 0.02* | *1.73 ± 0.24* | *0.83 ± 0.06* | *1.59 ± 0.20* |
| *Parietal* | *GM* | *R* | *Sample 2 (Geneva)* | *0.24 ± 0.02* | *1.52 ± 0.26* | *0.85 ± 0.11* | *1.57 ± 0.31* |
| *Parietal* | *WM* | *L* | *Sample 1 (Lausanne)* | *0.26 ± 0.02* | *1.55 ± 0.31* | *0.83 ± 0.05* | *1.84 ± 0.23* |
| *Parietal* | *WM* | *L* | *Sample 2 (Geneva)* | *0.28 ± 0.03* | *1.37 ± 0.28* | *0.86 ± 0.11* | *1.87 ± 0.30* |

| Brain region | Type | Side | Cohort | Cho / tCr | Glx / tCr | Ins / tCr | tNAA / tCr |
|---|---|---|---|---|---|---|---|
| *Parietal* | *WM* | *R* | *Sample 1 (Lausanne)* | *0.25 ± 0.02* | *1.58 ± 0.32* | *0.83 ± 0.05* | *1.80 ± 0.23* |
| *Parietal* | *WM* | *R* | *Sample 2 (Geneva)* | *0.28 ± 0.03* | *1.37 ± 0.28* | *0.86 ± 0.11* | *1.88 ± 0.30* |
| *Temporal* | *GM* | *L* | *Sample 1 (Lausanne)* | *0.26 ± 0.02* | *1.71 ± 0.29* | *0.84 ± 0.09* | *1.45 ± 0.23* |
| *Temporal* | *GM* | *L* | *Sample 2 (Geneva)* | *0.27 ± 0.04* | *1.50 ± 0.33* | *0.86 ± 0.14* | *1.41 ± 0.32* |
| *Temporal* | *GM* | *R* | *Sample 1 (Lausanne)* | *0.28 ± 0.06* | *1.71 ± 0.24* | *0.87 ± 0.14* | *1.37 ± 0.20* |
| *Temporal* | *GM* | *R* | *Sample 2 (Geneva)* | *0.28 ± 0.04* | *1.54 ± 0.36* | *0.87 ± 0.13* | *1.42 ± 0.33* |
| *Temporal* | *WM* | *L* | *Sample 1 (Lausanne)* | *0.29 ± 0.04* | *1.50 ± 0.22* | *0.91 ± 0.21* | *1.69 ± 0.21* |
| *Temporal* | *WM* | *L* | *Sample 2 (Geneva)* | *0.31 ± 0.04* | *1.37 ± 0.30* | *0.87 ± 0.14* | *1.70 ± 0.32* |
| *Temporal* | *WM* | *R* | *Sample 1 (Lausanne)* | *0.29 ± 0.03* | *1.53 ± 0.21* | *0.88 ± 0.09* | *1.63 ± 0.22* |
| *Temporal* | *WM* | *R* | *Sample 2 (Geneva)* | *0.31 ± 0.03* | *1.38 ± 0.30* | *0.88 ± 0.12* | *1.69 ± 0.33* |
| *Occipital* | *GM* | *L* | *Sample 1 (Lausanne)* | *0.21 ± 0.01* | *1.70 ± 0.31* | *0.82 ± 0.05* | *1.61 ± 0.20* |
| *Occipital* | *GM* | *L* | *Sample 2 (Geneva)* | *0.22 ± 0.03* | *1.46 ± 0.28* | *0.85 ± 0.12* | *1.57 ± 0.33* |
| *Occipital* | *GM* | *R* | *Sample 1 (Lausanne)* | *0.21 ± 0.02* | *1.73 ± 0.32* | *0.83 ± 0.05* | *1.58 ± 0.21* |
| *Occipital* | *GM* | *R* | *Sample 2 (Geneva)* | *0.22 ± 0.03* | *1.45 ± 0.26* | *0.85 ± 0.12* | *1.60 ± 0.34* |

| Brain region | Type | Side | Cohort | Cho / tCr | Glx / tCr | Ins / tCr | tNAA / tCr |
|---|---|---|---|---|---|---|---|
| *Occipital* | *WM* | *L* | *Sample 1 (Lausanne)* | *0.24 ± 0.01* | *1.57 ± 0.32* | *0.86 ± 0.06* | *1.81 ± 0.20* |
| *Occipital* | *WM* | *L* | *Sample 2 (Geneva)* | *0.25 ± 0.03* | *1.37 ± 0.28* | *0.86 ± 0.12* | *1.78 ± 0.33* |
| *Occipital* | *WM* | *R* | *Sample 1 (Lausanne)* | *0.23 ± 0.01* | *1.47 ± 0.22* | *0.85 ± 0.07* | *1.76 ± 0.19* |
| *Occipital* | *WM* | *R* | *Sample 2 (Geneva)* | *0.25 ± 0.03* | *1.33 ± 0.25* | *0.86 ± 0.11* | *1.78 ± 0.33* |
| *Thalamus* | *GM* | *L* | *Sample 1 (Lausanne)* | *0.35 ± 0.03* | *1.50 ± 0.33* | *0.82 ± 0.08* | *1.74 ± 0.25* |
| *Thalamus* | *GM* | *L* | *Sample 2 (Geneva)* | *0.35 ± 0.05* | *1.32 ± 0.33* | *0.81 ± 0.12* | *1.75 ± 0.33* |
| *Thalamus* | *GM* | *R* | *Sample 1 (Lausanne)* | *0.34 ± 0.02* | *1.58 ± 0.31* | *0.82 ± 0.07* | *1.75 ± 0.30* |
| *Thalamus* | *GM* | *R* | *Sample 2 (Geneva)* | *0.35 ± 0.05* | *1.31 ± 0.30* | *0.81 ± 0.12* | *1.75 ± 0.35* |
| *Putamen* | *GM* | *L* | *Sample 1 (Lausanne)* | *0.30 ± 0.04* | *1.41 ± 0.36* | *0.76 ± 0.08* | *1.54 ± 0.20* |
| *Putamen* | *GM* | *L* | *Sample 2 (Geneva)* | *0.30 ± 0.04* | *1.32 ± 0.35* | *0.76 ± 0.12* | *1.52 ± 0.31* |
| *Putamen* | *GM* | *R* | *Sample 1 (Lausanne)* | *0.29 ± 0.02* | *1.40 ± 0.31* | *0.73 ± 0.08* | *1.50 ± 0.27* |
| *Putamen* | *GM* | *R* | *Sample 2 (Geneva)* | *0.30 ± 0.04* | *1.29 ± 0.34* | *0.75 ± 0.11* | *1.57 ± 0.34* |
| *Pallidum* | *GM* | *L* | *Sample 1 (Lausanne)* | *0.29 ± 0.04* | *1.44 ± 0.46* | *0.72 ± 0.09* | *1.58 ± 0.26* |
| *Pallidum* | *GM* | *L* | *Sample 2 (Geneva)* | *0.29 ± 0.04* | *1.23 ± 0.33* | *0.71 ± 0.11* | *1.59 ± 0.34* |

| Brain region | Type | Side | Cohort | Cho / tCr | Glx / tCr | Ins / tCr | tNAA / tCr |
|---|---|---|---|---|---|---|---|
| *Pallidum* | *GM* | *R* | *Sample 1 (Lausanne)* | *0.28 ± 0.03* | *1.40 ± 0.45* | *0.70 ± 0.10* | *1.54 ± 0.28* |
| *Pallidum* | *GM* | *R* | *Sample 2 (Geneva)* | *0.29 ± 0.04* | *1.21 ± 0.34* | *0.70 ± 0.11* | *1.62 ± 0.35* |
| *Caudate* | *GM* | *L* | *Sample 1 (Lausanne)* | *0.36 ± 0.05* | *1.42 ± 0.34* | *0.89 ± 0.15* | *1.70 ± 0.24* |
| *Caudate* | *GM* | *L* | *Sample 2 (Geneva)* | *0.36 ± 0.06* | *1.36 ± 0.39* | *0.82 ± 0.15* | *1.61 ± 0.34* |
| *Caudate* | *GM* | *R* | *Sample 1 (Lausanne)* | *0.36 ± 0.04* | *1.47 ± 0.35* | *0.88 ± 0.15* | *1.76 ± 0.24* |
| *Caudate* | *GM* | *R* | *Sample 2 (Geneva)* | *0.36 ± 0.08* | *1.30 ± 0.37* | *0.84 ± 0.22* | *1.65 ± 0.38* |

#### Table S4. Cramer-Rao Lower Bound (CRLB) for each metabolite, by brain region

| Brain Region | Type | Side | Cohort | CRLB tCr | CRLB Cho | CRLB Glx | CRLB Ins | CRLB tNAA |
|---|---|---|---|---|---|---|---|---|
| *Frontal* | *GM* | *L* | *Sample 1 (Lausanne)* | *4.35 ± 1.20* | *5.53 ± 1.28* | *6.72 ± 2.11* | *5.90 ± 1.48* | *3.76 ± 1.24* |
| *Frontal* | *GM* | *L* | *Sample 2 (Geneva)* | *5.49 ± 3.76* | *6.38 ± 3.70* | *8.06 ± 3.30* | *6.64 ± 3.53* | *4.83 ± 3.11* |
| *Frontal* | *GM* | *R* | *Sample 1 (Lausanne)* | *4.38 ± 1.09* | *5.51 ± 1.15* | *6.73 ± 2.02* | *5.90 ± 1.34* | *3.80 ± 1.15* |
| *Frontal* | *GM* | *R* | *Sample 2 (Geneva)* | *5.71 ± 3.91* | *6.36 ± 2.34* | *8.68 ± 5.33* | *6.70 ± 3.10* | *4.95 ± 3.22* |
| *Frontal* | *WM* | *L* | *Sample 1 (Lausanne)* | *3.79 ± 0.76* | *4.69 ± 0.82* | *6.80 ± 1.96* | *5.36 ± 0.98* | *2.99 ± 0.72* |
| *Frontal* | *WM* | *L* | *Sample 2 (Geneva)* | *4.62 ± 2.47* | *5.30 ± 1.79* | *8.45 ± 3.34* | *6.14 ± 2.54* | *3.64 ± 1.66* |
| *Frontal* | *WM* | *R* | *Sample 1 (Lausanne)* | *3.75 ± 0.71* | *4.65 ± 0.81* | *6.65 ± 1.70* | *5.31 ± 0.92* | *2.98 ± 0.63* |
| *Frontal* | *WM* | *R* | *Sample 2 (Geneva)* | *4.73 ± 2.72* | *5.31 ± 1.53* | *8.90 ± 5.29* | *6.19 ± 2.02* | *3.77 ± 1.62* |
| *Parietal* | *GM* | *L* | *Sample 1 (Lausanne)* | *3.46 ± 1.18* | *4.87 ± 1.26* | *6.12 ± 2.27* | *5.07 ± 1.46* | *3.10 ± 1.11* |
| *Parietal* | *GM* | *L* | *Sample 2 (Geneva)* | *4.24 ± 2.32* | *5.49 ± 1.92* | *7.14 ± 2.66* | *5.69 ± 2.26* | *3.83 ± 2.18* |
| *Parietal* | *GM* | *R* | *Sample 1 (Lausanne)* | *3.54 ± 1.04* | *4.93 ± 1.21* | *5.98 ± 1.82* | *5.13 ± 1.35* | *3.19 ± 1.01* |
| *Parietal* | *GM* | *R* | *Sample 2 (Geneva)* | *4.41 ± 3.40* | *5.51 ± 1.81* | *7.43 ± 3.97* | *5.89 ± 3.60* | *3.59 ± 1.71* |
| *Parietal* | *WM* | *L* | *Sample 1 (Lausanne)* | *3.03 ± 0.80* | *4.22 ± 0.88* | *6.23 ± 1.96* | *4.66 ± 0.99* | *2.48 ± 0.67* |

| Brain Region | Type | Side | Cohort | CRLB tCr | CRLB Cho | CRLB Glx | CRLB Ins | CRLB tNAA |
|---|---|---|---|---|---|---|---|---|
| *Parietal* | *WM* | *L* | *Sample 2 (Geneva)* | *3.24 ± 0.73* | *4.42 ± 0.85* | *7.33 ± 2.34* | *5.03 ± 1.24* | *2.54 ± 0.65* |
| *Parietal* | *WM* | *R* | *Sample 1 (Lausanne)* | *3.08 ± 0.74* | *4.28 ± 0.87* | *6.03 ± 1.43* | *4.65 ± 0.87* | *2.52 ± 0.52* |
| *Parietal* | *WM* | *R* | *Sample 2 (Geneva)* | *3.32 ± 0.93* | *4.52 ± 0.97* | *7.52 ± 2.67* | *5.10 ± 1.34* | *2.62 ± 0.72* |
| *Temporal* | *GM* | *L* | *Sample 1 (Lausanne)* | *3.58 ± 0.91* | *4.95 ± 1.08* | *6.15 ± 1.60* | *5.25 ± 1.10* | *3.25 ± 0.90* |
| *Temporal* | *GM* | *L* | *Sample 2 (Geneva)* | *4.35 ± 3.28* | *5.30 ± 2.13* | *7.21 ± 3.32* | *5.64 ± 2.44* | *3.66 ± 2.62* |
| *Temporal* | *GM* | *R* | *Sample 1 (Lausanne)* | *3.77 ± 0.89* | *5.15 ± 1.07* | *6.19 ± 1.43* | *5.31 ± 1.03* | *3.36 ± 0.89* |
| *Temporal* | *GM* | *R* | *Sample 2 (Geneva)* | *4.53 ± 2.49* | *5.51 ± 1.73* | *7.81 ± 4.30* | *5.87 ± 2.56* | *3.72 ± 1.90* |
| *Temporal* | *WM* | *L* | *Sample 1 (Lausanne)* | *3.40 ± 0.72* | *4.44 ± 0.84* | *6.81 ± 1.47* | *5.09 ± 0.90* | *2.78 ± 0.61* |
| *Temporal* | *WM* | *L* | *Sample 2 (Geneva)* | *3.93 ± 1.38* | *4.77 ± 1.23* | *8.28 ± 3.66* | *5.92 ± 2.90* | *3.16 ± 1.07* |
| *Temporal* | *WM* | *R* | *Sample 1 (Lausanne)* | *3.47 ± 0.79* | *4.55 ± 0.90* | *6.73 ± 1.27* | *5.08 ± 0.94* | *2.91 ± 0.76* |
| *Temporal* | *WM* | *R* | *Sample 2 (Geneva)* | *4.07 ± 1.54* | *4.90 ± 1.36* | *8.24 ± 3.13* | *5.74 ± 1.95* | *3.26 ± 1.27* |
| *Occipital* | *GM* | *L* | *Sample 1 (Lausanne)* | *3.91 ± 1.25* | *5.62 ± 1.62* | *6.77 ± 1.80* | *5.59 ± 1.53* | *3.56 ± 1.21* |
| *Occipital* | *GM* | *L* | *Sample 2 (Geneva)* | *4.46 ± 3.51* | *6.21 ± 4.09* | *7.76 ± 3.92* | *6.00 ± 3.40* | *3.55 ± 1.74* |
| *Occipital* | *GM* | *R* | *Sample 1 (Lausanne)* | *4.00 ± 1.44* | *5.81 ± 1.86* | *6.76 ± 1.93* | *5.61 ± 1.67* | *3.61 ± 1.43* |

| Brain Region | Type | Side | Cohort | CRLB tCr | CRLB Cho | CRLB Glx | CRLB Ins | CRLB tNAA |
|---|---|---|---|---|---|---|---|---|
| *Occipital* | *GM* | *R* | *Sample 2 (Geneva)* | *4.56 ± 3.25* | *6.22 ± 3.22* | *8.26 ± 4.93* | *6.15 ± 3.37* | *3.95 ± 3.02* |
| *Occipital* | *WM* | *L* | *Sample 1 (Lausanne)* | *3.56 ± 0.99* | *4.97 ± 1.28* | *7.00 ± 1.54* | *5.15 ± 1.23* | *2.93 ± 0.83* |
| *Occipital* | *WM* | *L* | *Sample 2 (Geneva)* | *4.15 ± 3.86* | *5.55 ± 3.83* | *8.61 ± 5.27* | *5.81 ± 3.43* | *3.12 ± 1.76* |
| *Occipital* | *WM* | *R* | *Sample 1 (Lausanne)* | *3.70 ± 1.18* | *5.19 ± 1.45* | *7.03 ± 1.62* | *5.30 ± 1.39* | *3.09 ± 1.13* |
| *Occipital* | *WM* | *R* | *Sample 2 (Geneva)* | *3.75 ± 1.42* | *5.04 ± 1.27* | *8.77 ± 6.12* | *5.32 ± 1.71* | *2.99 ± 1.21* |
| *Thalamus* | *GM* | *L* | *Sample 1 (Lausanne)* | *4.81 ± 1.08* | *5.08 ± 0.93* | *8.48 ± 2.95* | *6.74 ± 1.10* | *3.87 ± 1.00* |
| *Thalamus* | *GM* | *L* | *Sample 2 (Geneva)* | *5.72 ± 10.39* | *6.29 ± 12.84* | *9.83 ± 5.53* | *6.84 ± 4.25* | *3.63 ± 1.55* |
| *Thalamus* | *GM* | *R* | *Sample 1 (Lausanne)* | *4.90 ± 1.29* | *5.23 ± 1.23* | *8.27 ± 2.31* | *6.83 ± 1.31* | *4.02 ± 1.26* |
| *Thalamus* | *GM* | *R* | *Sample 2 (Geneva)* | *5.27 ± 8.87* | *6.00 ± 10.09* | *9.79 ± 5.06* | *7.10 ± 6.98* | *3.72 ± 2.14* |
| *Putamen* | *GM* | *L* | *Sample 1 (Lausanne)* | *4.48 ± 0.87* | *5.16 ± 0.87* | *8.97 ± 3.22* | *6.49 ± 1.12* | *3.80 ± 0.89* |
| *Putamen* | *GM* | *L* | *Sample 2 (Geneva)* | *4.85 ± 2.64* | *5.96 ± 3.68* | *9.42 ± 4.55* | *7.14 ± 3.05* | *4.59 ± 3.81* |
| *Putamen* | *GM* | *R* | *Sample 1 (Lausanne)* | *4.12 ± 0.72* | *4.93 ± 0.80* | *7.79 ± 2.48* | *6.43 ± 1.07* | *3.43 ± 0.82* |
| *Putamen* | *GM* | *R* | *Sample 2 (Geneva)* | *4.59 ± 2.71* | *5.63 ± 2.61* | *9.16 ± 3.23* | *6.76 ± 2.14* | *4.23 ± 2.90* |
| *Pallidum* | *GM* | *L* | *Sample 1 (Lausanne)* | *5.13 ± 1.54* | *6.22 ± 1.79* | *10.13 ± 4.10* | *7.72 ± 2.09* | *4.40 ± 1.70* |

| **Brain Region** | **Type** | **Side** | **Cohort** | **CRLB tCr** | **CRLB Cho** | **CRLB Glx** | **CRLB Ins** | **CRLB tNAA** |
|---|---|---|---|---|---|---|---|---|
| *Pallidum* | *GM* | *L* | *Sample 2 (Geneva)* | *9.51 ± 21.05* | *6.52 ± 3.60* | *10.85 ± 5.79* | *9.13 ± 9.37* | *6.08 ± 11.28* |
| *Pallidum* | *GM* | *R* | *Sample 1 (Lausanne)* | *5.28 ± 1.49* | *6.18 ± 1.41* | *9.10 ± 2.83* | *7.60 ± 1.73* | *4.28 ± 1.24* |
| *Pallidum* | *GM* | *R* | *Sample 2 (Geneva)* | *6.73 ± 11.15* | *5.99 ± 1.82* | *10.39 ± 3.63* | *7.71 ± 2.27* | *4.70 ± 3.23* |
| *Caudate* | *GM* | *L* | *Sample 1 (Lausanne)* | *5.04 ± 1.08* | *5.41 ± 1.12* | *9.22 ± 2.69* | *6.64 ± 1.27* | *3.88 ± 0.93* |
| *Caudate* | *GM* | *L* | *Sample 2 (Geneva)* | *7.80 ± 13.17* | *6.14 ± 3.39* | *11.08 ± 5.92* | *8.38 ± 7.04* | *5.90 ± 9.67* |
| *Caudate* | *GM* | *R* | *Sample 1 (Lausanne)* | *4.65 ± 1.14* | *4.97 ± 0.99* | *8.25 ± 2.58* | *6.45 ± 1.01* | *3.76 ± 1.01* |
| *Caudate* | *GM* | *R* | *Sample 2 (Geneva)* | *6.73 ± 13.58* | *5.77 ± 2.45* | *10.83 ± 7.79* | *7.20 ± 2.79* | *4.52 ± 2.17* |

#### Table S5. Full Width at Half Maximum (FWHM) and Signal-to-Noise Ratio (SNR), by brain region

| Brain Region | Type | Side | Cohort | FWHM | SNR |
|---|---|---|---|---|---|
| *Frontal* | *GM* | *L* | *Sample 1 (Lausanne)* | *0.05 ± 0.00* | *15.96 ± 5.48* |
| *Frontal* | *GM* | *L* | *Sample 2 (Geneva)* | *0.05 ± 0.01* | *15.88 ± 4.79* |
| *Frontal* | *GM* | *R* | *Sample 1 (Lausanne)* | *0.05 ± 0.00* | *15.73 ± 5.30* |
| *Frontal* | *GM* | *R* | *Sample 2 (Geneva)* | *0.05 ± 0.01* | *15.36 ± 4.76* |
| *Frontal* | *WM* | *L* | *Sample 1 (Lausanne)* | *0.04 ± 0.00* | *19.42 ± 4.72* |
| *Frontal* | *WM* | *L* | *Sample 2 (Geneva)* | *0.04 ± 0.00* | *18.88 ± 4.40* |
| *Frontal* | *WM* | *R* | *Sample 1 (Lausanne)* | *0.04 ± 0.00* | *19.52 ± 4.50* |
| *Frontal* | *WM* | *R* | *Sample 2 (Geneva)* | *0.04 ± 0.00* | *18.65 ± 4.39* |
| *Parietal* | *GM* | *L* | *Sample 1 (Lausanne)* | *0.04 ± 0.00* | *21.44 ± 7.23* |
| *Parietal* | *GM* | *L* | *Sample 2 (Geneva)* | *0.04 ± 0.00* | *19.99 ± 5.89* |
| *Parietal* | *GM* | *R* | *Sample 1 (Lausanne)* | *0.04 ± 0.00* | *20.97 ± 6.79* |
| *Parietal* | *GM* | *R* | *Sample 2 (Geneva)* | *0.04 ± 0.00* | *20.09 ± 6.19* |
| *Parietal* | *WM* | *L* | *Sample 1 (Lausanne)* | *0.04 ± 0.00* | *24.62 ± 6.34* |
| *Parietal* | *WM* | *L* | *Sample 2 (Geneva)* | *0.03 ± 0.00* | *23.69 ± 5.61* |
| *Parietal* | *WM* | *R* | *Sample 1 (Lausanne)* | *0.04 ± 0.00* | *24.58 ± 5.96* |
| *Parietal* | *WM* | *R* | *Sample 2 (Geneva)* | *0.03 ± 0.00* | *23.59 ± 5.82* |
| *Temporal* | *GM* | *L* | *Sample 1 (Lausanne)* | *0.05 ± 0.00* | *15.44 ± 5.01* |
| *Temporal* | *GM* | *L* | *Sample 2 (Geneva)* | *0.04 ± 0.01* | *15.62 ± 4.64* |
| *Temporal* | *GM* | *R* | *Sample 1 (Lausanne)* | *0.05 ± 0.00* | *14.96 ± 4.46* |
| *Temporal* | *GM* | *R* | *Sample 2 (Geneva)* | *0.04 ± 0.01* | *15.18 ± 4.29* |
| *Temporal* | *WM* | *L* | *Sample 1 (Lausanne)* | *0.04 ± 0.00* | *18.28 ± 4.62* |
| *Temporal* | *WM* | *L* | *Sample 2 (Geneva)* | *0.03 ± 0.00* | *18.21 ± 4.81* |

| Brain Region | Type | Side | Cohort | FWHM | SNR |
|---|---|---|---|---|---|
| *Temporal* | *WM* | *R* | *Sample 1 (Lausanne)* | *0.04 ± 0.00* | *17.92 ± 4.41* |
| *Temporal* | *WM* | *R* | *Sample 2 (Geneva)* | *0.04 ± 0.00* | *17.77 ± 4.49* |
| *Occipital* | *GM* | *L* | *Sample 1 (Lausanne)* | *0.04 ± 0.00* | *19.29 ± 7.18* |
| *Occipital* | *GM* | *L* | *Sample 2 (Geneva)* | *0.04 ± 0.00* | *19.02 ± 6.22* |
| *Occipital* | *GM* | *R* | *Sample 1 (Lausanne)* | *0.04 ± 0.00* | *18.72 ± 7.02* |
| *Occipital* | *GM* | *R* | *Sample 2 (Geneva)* | *0.04 ± 0.00* | *19.29 ± 6.02* |
| *Occipital* | *WM* | *L* | *Sample 1 (Lausanne)* | *0.04 ± 0.00* | *21.41 ± 6.57* |
| *Occipital* | *WM* | *L* | *Sample 2 (Geneva)* | *0.03 ± 0.00* | *21.42 ± 6.42* |
| *Occipital* | *WM* | *R* | *Sample 1 (Lausanne)* | *0.04 ± 0.00* | *21.17 ± 6.68* |
| *Occipital* | *WM* | *R* | *Sample 2 (Geneva)* | *0.03 ± 0.00* | *21.49 ± 6.19* |
| *Thalamus* | *GM* | *L* | *Sample 1 (Lausanne)* | *0.04 ± 0.00* | *14.10 ± 2.82* |
| *Thalamus* | *GM* | *L* | *Sample 2 (Geneva)* | *0.04 ± 0.00* | *16.21 ± 4.25* |
| *Thalamus* | *GM* | *R* | *Sample 1 (Lausanne)* | *0.04 ± 0.00* | *14.44 ± 3.16* |
| *Thalamus* | *GM* | *R* | *Sample 2 (Geneva)* | *0.04 ± 0.00* | *16.04 ± 3.92* |
| *Putamen* | *GM* | *L* | *Sample 1 (Lausanne)* | *0.05 ± 0.01* | *13.46 ± 3.03* |
| *Putamen* | *GM* | *L* | *Sample 2 (Geneva)* | *0.04 ± 0.01* | *14.29 ± 3.87* |
| *Putamen* | *GM* | *R* | *Sample 1 (Lausanne)* | *0.05 ± 0.01* | *14.34 ± 3.76* |
| *Putamen* | *GM* | *R* | *Sample 2 (Geneva)* | *0.04 ± 0.01* | *14.68 ± 3.73* |
| *Pallidum* | *GM* | *L* | *Sample 1 (Lausanne)* | *0.05 ± 0.01* | *11.65 ± 3.28* |
| *Pallidum* | *GM* | *L* | *Sample 2 (Geneva)* | *0.04 ± 0.01* | *12.88 ± 3.84* |
| *Pallidum* | *GM* | *R* | *Sample 1 (Lausanne)* | *0.05 ± 0.01* | *12.81 ± 4.48* |
| *Pallidum* | *GM* | *R* | *Sample 2 (Geneva)* | *0.04 ± 0.01* | *13.42 ± 4.09* |
| *Caudate* | *GM* | *L* | *Sample 1 (Lausanne)* | *0.05 ± 0.01* | *12.58 ± 2.85* |
| *Caudate* | *GM* | *L* | *Sample 2 (Geneva)* | *0.04 ± 0.01* | *12.93 ± 3.35* |

| **Brain Region** | **Type** | **Side** | **Cohort** | **FWHM** | **SNR** |
|---|---|---|---|---|---|
| *Caudate* | *GM* | *R* | *Sample 1 (Lausanne)* | *0.04 ± 0.01* | *13.82 ± 3.43* |
| *Caudate* | *GM* | *R* | *Sample 2 (Geneva)* | *0.04 ± 0.01* | *13.32 ± 3.04* |

#### Table S6. MRSinMRS informations

| Category | Geneva Study | Lausanne Psychosis Cohort |
|---|---|---|
| Scanner | 3T Magnetom TrioTim (Siemens) | 3T Prisma Fit (Siemens) |
| RF coils | 32 ch 1H head coil | 32 ch 1H head coil |
| Coil elements | HEA;HEP | HEA;HEP |
| Sequence | 3D 1H-FID-MRSI (CS-accelerated) | 3D 1H-FID-MRSI (CS-accelerated) |
| Position | R4.8 A11.8 H36.9 | L3.9 A23.0 H4.3 |
| Orientation | T > C-13.5 > S2.5 | T > C34.5 > S-5.4 |
| Rotation (deg) | -2 | 0 |
| TE (ms) | 1.5 | 1.0 |
| TR (ms) | 372 | 353 |
| Averages | 1 | 1 |
| Flip angle (°) | 35 | 40 |
| FOV (mm) | 210 × 160 × 105 | 210 × 160 × 95 |
| Slab thickness (mm) | 95 | 95 |
| Slabs | 1 | 1 |
| Resolution ($mm^3$) | 5 × 5 × 5.3 | 5 × 5 × 5.3 |
| Spectral bandwidth (Hz) | 2000 | 2000 |
| FID points / Vector size | 512 | 512 |
| Acquisition duration (ms) | 256 | 256 |
| Matrix size | 32 × 42 × 20 | 32 × 42 × 20 |
| Water reference TE (ms) | 1.5 | 1.07 |

| Category | Geneva Study | Lausanne Psychosis Cohort |
|---|---|---|
| Water reference TR (ms) | 36 | 25 |
| Water reference flip angle (°) | 3 | 5 |
| Water reference resolution ($mm^3$) | 6.6 × 6.7 × 6.6 | 6.6 × 6.7 × 6.6 |
| Water reference FID points | 16 | 16 |
| Averaging mode | Short term | Short term |
| Water suppr. | Weak water suppr. | Water sat. |
| Water suppr. BW (Hz) | 60 | 60 |
| Spectral suppr. | None | None |
| Measurements | 1 | 1 |
| Saturation bands | 2 bands, 20 mm thickness | 2 bands, 20 mm thickness |
| Compressed sensing | acceleration factor 3.3, radius 0.2 | acceleration factor 3.3, radius 0.2 |
| Preparation scans | 4 | 4 |
| Dimension | 3D | 3D |
| Delta frequency (ppm) | 0.00 | 0.00 |
| Phase encoding | Elliptical | Elliptical |
| Remove oversampling | On | On |
| WS timing (ms) | 24 | 22 |
| WS amplitude factor | 0.9 | 0.9 |
| Max gradient amplitude used (mT/m) | 26.0 | 33.00 |
| Shim mode | Advanced | Advanced |

| Category | Geneva Study | Lausanne Psychosis Cohort |
|---|---|---|
| Data processing | Low-rank + TGV reconstruction; lipid/water removal | Low-rank + TGV reconstruction; lipid/water removal |
| Quantification | LCModel | LCModel |
| Metabolite basis set (LCModel) | NAA, NAAG, Cr, PCr, GPC, PCh, mI, sI, Glu, Gln, Lac, GABA, GSH, Tau, Asp, Ala | NAA, NAAG, Cr, PCr, GPC, PCh, mI, sI, Glu, Gln, Lac, GABA, GSH, Tau, Asp, Ala |
| Combined metabolites | tNAA (NAA+NAAG), tCr (Cr+PCr), Cho (GPC+PCh), Ins (mI), Glx (Glu+Gln) | tNAA (NAA+NAAG), tCr (Cr+PCr), Cho (GPC+PCh), Ins (mI), Glx (Glu+Gln) |
| Quality metrics | SNR, CRLB, FWHM | SNR, CRLB, FWHM |

#### Table S7: Control file of LCModel

```
$LCMODL
sddegz=999.
sddegp= 2.
degzer= 0.00
degppm= 0.00
ppmst= 4.3
ppmend= 1
nunfil= 2048
ndslic= 1
ndrows= 1
ndcols= 1
ltable= 7
lps= 8
lcoord= 9
islice= 1
irowst=1
irowen= 1
icolst= 1
icolen= 1
hzpppm= 123.1887
echot= 1500 (Geneva)| echot= 1 (Lausanne)
dows= T
nsimul= 0
dkntmn = 0.075
$END
```

### Table S8: Pre-processing and VBA analyses parameters sum-up

| Pre-processing and VBA analyses parameters sum-up | | |
|---|---|---|
| **Category** | **Parameter** | **Specification** |
| **Preprocessing** | Software | ANTs (v 2.6.2), PETPVC (v 1.2.12), FSL *bet2* (v6.0.7.7), cat12 (v12.9) |
| | Modality | Whole-brain metabolic maps (tNAA, Ins, Cho, Glx, tCr) |
| | Spatial resolution after normalization | $1 \times 1 \times 1$ $mm^3$ |
| | PVE correction | Region-based voxel-wise correction (PETPVC toolbox) |
| | Spike/outlier handling | $99^{th}$ percentile threshold; biharmonic inpainting + 3×3×3 median filtering |
| | Brain masking | Subject-level Qmask based on SNR >4, FWHM >0.1 or CRLB >20 ; Spikes also added to Qmask |
| | Registration (MRSI → T1w) | Rigid + SyN (cross-correlation metric), linear interpolation |
| | Registration (T1w → MNI) | Rigid + affine (mutual information) + SyN (cross-correlation), linear interpolation |
| | Interpolation order | Linear (for all transformations) |
| **Statistical model (GLM)** | Software | FSL *randomise* (v6.0.7.7) |
| | Model type | General linear model (between-group comparisons or correlations) |
| | Covariates | Age, sex for each analysis |
| | Group coding | Dummy-coded contrasts with mirror column for group comparison |
| | Voxel-wise exclusion of low-quality voxels | Integrated via *–mask* and lesion-masking approach (Winkler et al., 2014) |
| **Permutation testing** | Number of permutations | 10,000 |
| | Exchangeability blocks | None (cross-sectional design) |
| **Inference & correction** | Multiple comparison correction | TFCE (Threshold-Free Cluster Enhancement) |
| | TFCE parameters | FSL defaults (E=0.5, H=2.0, connectivity=6) |
| | Significance threshold | $p < 0.05$, family-wise error corrected |

| | | |
|---|---|---|
| | Reporting | Significant clusters displayed in MNI152 space; mean values extracted for post-hoc tests |
| **Masking / search space** | Gray/white matter restriction | Whole-brain GM and WM (for metabolic VBA) |
| | WM-only mask (for gFA) | FMRIB58_FA_1mm, threshold 0.25 |
| **Quality control** | Visual QC | Registration and normalization checked for all subjects |
| | Automated QC thresholds | CRLB < 20; FWHM < 0.1; SNR > 4 |

### Table S9: Coefficient of variation (COV) of tNAA compared to tCr in patients and controls

| | **tNAA** | **tCr** | **Difference** (95%CI) | *p-value* |
|---|---|---|---|---|
| Patients | 18.2% | 13.7% | 0.030-0.058 | 0.470 |
| Controls | 17.3% | 15.2% | 0.001-0.041 | 0.494 |
